# Predicting individual speech intelligibility from the cortical tracking of acoustic- and phonetic-level speech representations

**DOI:** 10.1101/471367

**Authors:** D Lesenfants, J Vanthornhout, E Verschueren, L Decruy, T Francart

**Author notes:** **Corresponding Author:** Tom Francart, Ph.D., KU Leuven Department of Neurosciences, ExpORL, Herestraat 49 bus 721, B-3000 Leuven, Belgium.

## Abstract

**Objective:** To objectively measure speech intelligibility of individual subjects from the EEG, based on cortical tracking of different representations of speech: low-level acoustical, higher-level discrete, or a combination. To compare each model’s prediction of the speech reception threshold (SRT) for each individual with the behaviorally measured SRT.

**Methods:** Nineteen participants listened to Flemish Matrix sentences presented at different signal-to-noise ratios (SNRs), corresponding to different levels of speech understanding. For different EEG frequency bands (delta, theta, alpha, beta or low-gamma), a model was built to predict the EEG signal from various speech representations: envelope, spectrogram, phonemes, phonetic features or a combination of phonetic Features and Spectrogram (FS). The same model was used for all subjects. The model predictions were then compared to the actual EEG of each subject for the different SNRs, and the prediction accuracy in function of SNR was used to predict the SRT.

**Results:** The model based on the FS speech representation and the theta EEG band yielded the best SRT predictions, with a difference between the behavioral and objective SRT below 1 decibel for 53% and below 2 decibels for 89% of the subjects.

**Conclusion:** A model including low- and higher-level speech features allows to predict the speech reception threshold from the EEG of people listening to natural speech. It has potential applications in diagnostics of the auditory system.

**Search Terms:** cortical speech tracking, objective measure, speech intelligibility, auditory processing, speech representations.

**Highlights:**

- Objective EEG-based measure of speech intelligibility
- Improved prediction of speech intelligibility by combining speech representations
- Cortical tracking of speech in the delta EEG band monotonically increased with SNRs
- Cortical responses in the theta EEG band best predicted the speech reception threshold

**Disclosure:** The authors report no disclosures relevant to the manuscript.

## INTRODUCTION

Nearly six percent of the world population has a disabling hearing loss, i.e., hearing loss greater than 40 decibels (dB) in the better hearing ear (report of the World Health Organization 2013). It is estimated that by 2060 this prevalence will double (Goman et al. 2017). Untreated hearing loss has been linked to cognitive decline, anxiety, depression, and social exclusion (Li et al. 2014), highlighting the importance of adequately evaluating and treating hearing disabilities. Current clinical measurement of hearing impairment relies on behavioral evaluation of hearing. While this gold standard is relatively fast and reliable in healthy adults, when performed by a well-trained audiologist with suitable equipment, it requires manual intervention, which is labor intensive, subjective and depends on the examiner’s experience and patient’s active participation. Moreover, the patient’s engagement can fluctuate over time (e.g., children) or even be absent (e.g., unconscious patients, see Lesenfants et al. 2016).

Electroencephalography (EEG)-based measures of hearing might provide a good alternative, as they are objective and can be performed automatically. However, current clinically-available objective measures of hearing, based on auditory brainstem responses (Jewett et al. 1970; Verhaert et al. 2008; Anderson et al. 2013) or auditory steady state responses (Galambos et al. 1981; Stapells et al. 1984; Picton et al. 2005; Luts et al. 2006; Gransier et al. 2016), can only complement behavioral evaluation. This is mainly because they rely on simple non-realistic auditory stimuli (e.g. clicks) leading to a measure of hearing but not speech understanding. A measure of brain activity in response to natural running speech could overcome these issues and provide a realistic and objective measure of a patient’s speech intelligibility (i.e., how well the person can understand speech) in clinical routine.

Because the temporal fluctuations of the speech stimulus envelope are critical for speech understanding (Shannon et al. 1995), in particular modulation frequencies below 20 Hz (Drullman et al. 1994b; Drullman et al. 1994a), many researchers (Aiken & Picton 2008; Ding & Simon 2013; Gonçalves et al. 2014; Biesmans et al. 2017) have evaluated the potential of measuring cortical tracking of the speech envelope using surface EEG. Recently, Vanthornhout et al. (2018) proposed to objectively measure speech intelligibility using cortical tracking of the speech envelope and showed a strong correlation between a behavioral and a novel electrophysiological measure of speech intelligibility. However, while these results are promising, the correlation between the behavioral and objective measures of speech intelligibility only explained 50% of the variance, and the objective measure could only be derived in three-quarter of the subjects. Recently, Di Liberto et al. (2015, 2016 & 2017) showed that the cortical tracking of running speech is better characterized using a model integrating both low-level spectro-temporal speech information (e.g., the speech envelope) and discrete higher-level phonetic features.

We here aimed to evaluate the potential of using speech representations beyond the envelope to objectively measure speech intelligibility. We constructed models based on different speech representations and evaluated their performance in different frequency bands. The key outcome measure was the difference between the gold standard speech reception threshold (SRT; the SNR at which a person can correctly recall 50% of spoken words), and the SRT predicted from the EEG. Stimulus representations beyond the envelope on the one hand allow to measure higher order neural processes, more closely related to speech intelligibility, and on the other hand allow higher EEG prediction accuracies. Therefore, following Di Liberto and colleagues, we hypothesized that the SRT could be better predicted using a more complex model integrating both acoustic and phonetic representations of the stimulus.

## METHODS

We analyzed data acquired in our previous study aiming to predict speech intelligibility using cortical entrainment of the speech envelope with a linear backward model (Vanthornhout et al. 2018).

### Participants

Nineteen native Flemish-speaking volunteers (7 men; age 24 ± 2 years; 2 were left-handed) participated in this study. Each participant reported normal hearing, verified by pure tone audiometry (pure tone thresholds lower than 20 dB HL for 125 Hz to 8000 Hz using a MADSEN Orbiter 922–2 audiometer). The study was approved by the Medical Ethics Committee Research UZ KU Leuven/Research (KU Leuven, Belgium) with reference S59040 and all participants provided informed consent.

### Experiments

The experiments were conducted on a laptop running Windows using the APEX 3 (version 3.1) software platform developed at ExpORL (Dept. Neurosciences, KU Leuven) (Francart et al. 2008), an RME Multiface II sound card (RME, Haimhausen, Germany), and Etymotic ER-3A insert phones (Etymotic Research, Inc., IL, USA) which were electromagnetically shielded using CFL2 boxes from Perancea Ltd. (London, UK). The setup was calibrated in a 2-cm^3^ coupler (Brüel & Kjaer, type 4152, Nærum, Denmark) using the stationary speech weighted noise corresponding to the Matrix speech material. The experiments took place in an electromagnetically shielded and soundproofed room.

#### Behavioral

Each participant started the experiment with a behavioral evaluation of speech understanding using the Flemish Matrix sentences (Luts et al. 2014) presented at three fixed SNRs (−9.5; −6.5 and −3.5 dB SNR) around the SRT. For each SNR, we counted the number of correctly repeated words following the presentation of each of 20 Matrix sentences. Then, we fitted a psychometric function through the data points and used its 50%-correct point as the behavioral SRT. This method is currently the gold standard in measuring speech intelligibility, both in research and clinical practice (Jansen et al. 2012; Decruy, Das, et al. 2018). The approximate total duration of the behavioral experiment was 30 minutes (including breaks at the discretion of the participant).

Sentences were spoken by a female speaker and presented to the right ear. Each sentence of the Flemish Matrix material is composed of 5 words that follow a fixed syntactic structure of *Name* + *Verb* + *Numeral* + *Adjective* + *Object* (e.g., “Sofie ziet zes grijze pennen”; “Sofie sees six gray pens”), each of them randomly selected from a set of 10 alternatives, each combination yielding similar behavioral speech intelligibility scores. These sentences sound perfectly natural, but are grammatically trivial and completely semantically unpredictable, therefore minimizing the effect of higher order language processing on the result.

#### Electrophysiological

Each subject listened to the children’s story “Milan” (we will call this condition “Story”), written and narrated in Flemish by Stijn Vranken. The stimulus was 14 minutes long and was presented binaurally without any noise. We then presented binaurally sequences of 40 Flemish Matrix sentences, each lasting 2 min, at different SNRs, in random order, with the speech level fixed at 60 dBA. Silences between sentences ranged in duration between 0.8 and 1.2s. At the end of each 2-min stimulus, we asked the participant questions to ensure a satisfactory level of attention on the task (e.g. ‘How many times did you hear “gray pens”?’). Group 1 underwent four presentations of five different SNRs (−9.5, −7.6, −5.5, −1 and 100 dB SNR). Group 2 underwent three presentations of seven different SNRs (−12.5, −9.5, −6.5, −3.5, −0.5, 2.5 and 100 dB SNR). Note that “100 dB SNR” corresponds to “no noise” (hereinafter named quiet condition). The approximate duration of the electrophysiological experiment was 2 hours (including breaks at the discretion of the participant).

In both the behavioral and electrophysiological parts of our study, the noise added to the Matrix sentences was a stationary speech weighted noise with the same long-term average speech spectrum as the speech, ensuring maximal energetic masking.

### Recordings

EEG signals were recorded from 64 Ag/AgCl ring electrodes at a sampling frequency of 8192 Hz using a Biosemi ActiveTwo system (Amsterdam, Netherlands). The electrodes were placed on the scalp according to international 10-20 standards.

### Data analysis

All analyses were done with custom-made Matlab (R2016b) scripts and the mTRF Toolbox (Crosse et al. 2016; Gonçalves et al. 2014).

#### Speech features

We extracted five different representations of the speech stimulus, selected according to Di Liberto et al. (2015):

1. The broadband amplitude envelope (Env) was extracted as the absolute value of the (complex) Hilbert transform of the speech signal.
2. The spectrogram (Sgram) was obtained by first filtering the speech stimulus into 16 logarithmically-spaced speech frequency bands using zero phase Butterworth filters with 80 dB attenuation at 10 % outside the passband between 250 Hz and 8 kHz, according to Greenwood’s equation (Greenwood 1961), assuming linear spacing in the cochlea. We then calculated the energy in each frequency band using a Hilbert transform (see Env).
3. The time-aligned sequence of phonemes (Ph) was extracted using the speech alignment component of the reading tutor (Duchateau et al. 2009), which allows for reading miscues (skipped, repeated, misread words), automatically segmenting each word into phonemes from the Dutch International Phonetic Alphabet (IPA) and performing forced alignment. We then converted this into a binary matrix mask representing the start and end time-points (i.e., a ‘1’ from the start until the end of each phoneme) for the 37 phonemes present in both the story and the matrix sentences. Two audiology/speech-language-therapy experts randomly selected six 2s-segments of the Matrix sentences and manually labelled them. The results of their labelling can be found in table Ap1 in the Appendix. Interestingly, the automatically extracted phoneme labels were 100% correct with a timing difference (i.e., the absolute difference in start/stop time) of only 11 ± 18 ms (median ± iqr).
4. The time-aligned binary sequence of phonetic features (Fea) was assembled using the following groups of phonemes: short vowels, long vowels, fricative consonants, nasal consonants and plosive consonants. Note that we decided to decrease the number of phonetic feature categories as compared to Di Liberto et al (2015), to ensure there were enough representations of each phonetic feature in each 2-minutes Matrix stimuli.
5. The combination of time-aligned sequence of phonetic Features and the Spectrogram (FS) as proposed by Di Liberto et al. (2015).

#### EEG signal processing

EEG signals were first downsampled from 8192 to 1024 Hz (using anti-aliasing filtering) to decrease processing time. EEG artifacts were then removed using a multi-channel Wiener filter algorithm with channel lags of three samples (i.e., all delays from −3 to 3 were included) and a noise-weighting factor of 1 (Somers et al. 2018). We then re-referenced each EEG signal to a common-average reference. EEG signals were bandpass filtered between 0.5-4 Hz (delta), 4-8 Hz (theta), 8-15 Hz (alpha), 15-30 Hz (beta), or 30-45 Hz (low gamma; for this frequency, we then computed the envelope of the filtered signals) using zero phase Butterworth filters with 80 dB attenuation at 10 % outside the passband. Stimulus representations and EEG signals were then further downsampled to 128 Hz (using anti-aliasing filtering). The impulse and step responses of the entire pipeline are shown in the Appendix (see Fig. Ap11), see also de Cheveigné & Nelken (2019). Note that while we needed to use multiple filters, the impact of filter artifacts on our main results should be minimal, as our main outcome measure is based on correlation between the actual and predicted EEG signal, which, contrary to direct assessment of temporal response functions is quite robust.

#### Temporal response function (TRF) estimation

For each subject, we first normalized both the speech representation (i.e., the Env or the Sgram; not the Ph or Fea since they consisted of a binary representation of the speech) and each EEG signal of the different conditions. A grand-average quantitative mapping between each speech representation and the EEG for the Story condition averaged across subjects (by averaging the covariance matrices of the EEG signals, Biesmans et al. 2017) was computed using ridge regression with a time-lag window of 0-400 ms. A visualization of each model’s TRF can be seen in Lesenfants et al. (2019). This grand-average mapping (based on the Story condition) was then used to predict EEG signals from each stimulus representation for the Matrix condition. Note that the clean speech features (without added noise) were used to predict the EEG signal, even for conditions in which noisy speech was presented to the subject, the rationale being that to understand speech the brain requires a de-noised version of the speech. Di Liberto et al (2017) showed that a single-subject model required at least 30-min of data in order to efficiently model both the low-level and higher-level speech features within a FS model, while 10-min of data per individual are enough when using a grand-average model. Therefore, given the intended clinical application of our method we decided to use a grand-average decoder in order to decrease the overall session time. The cortical tracking of speech was computed as the correlation (Spearman’s rank-order correlation) between the recorded and predicted EEG signals at the different electrode locations, for each subject. Note that with our grand average model, the predicted EEG is the same for each subject, as it only depends on the stimulus, but the actual EEG of course differs across subjects.

#### Models’ performance

For each model, we evaluated the extent to which we could predict SRT from the EEG. We will call the predicted SRT the correlation threshold (CT). Similar to Vanthornhout et al. (2018) inspired by the way the SRT is derived from the behavioral measurements, we fitted a sigmoid S to the Spearman’s correlation between actual and predicted EEG in function of SNR in order to obtain the CT using the formula:

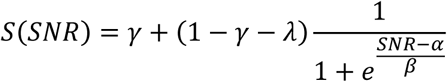

With γ the guess-rate, λ the lapse-rate, α the midpoint, and β the slope. Each subject’s CT was then computed as the averaged α across presentations (i.e., four presentations for group 1, three presentations for group 2), with only α in the range of −4 to −11 dB SNR taken into account in the averaging. If none of the α were in this range, CT was considered as non-assessed (NA, see Fig. 4).

#### Statistical analysis

A permutation test (Nichols & Holmes 2002; Noirhomme et al. 2014) was used to evaluate the significance level for each model (1000 repetitions, p < .01): for all electrode locations, we first predicted the EEG response to a speech feature using the trained model, we randomly permuted the predicted EEG samples 1000 times and calculated the Spearman’s correlation between the result and the actual EEG signal for each permutation. From the resulting distribution of correlations, the significance level was calculated as its 1^st^ percentile. The significance of change between conditions was assessed with a non-parametric Wilcoxon signed-rank test (2-tailed, p < .01), with Bonferroni correction for multiple comparisons.

## RESULTS

### Behavioral evaluation of SRT

The mean of the individual behavioral SRTs was −7.1 dB with an inter-subject standard deviation of 1.5 dB, ranging from −9.3 to −4.7 dB for group 1, and −8.5 dB with an inter-subject standard deviation of 0.8 dB, ranging from −10.3 to −7.7 dB for group 2.

### Spatial distribution of the cortical tracking

We calculated the cortical tracking over the scalp for the different EEG frequency bands averaged across the participants for the FS-model. Averaged cortical speech tracking in the delta EEG band increased with SNR until reaching a maximum in quiet for both group 1 (see Fig.1, row 1) and group 2 (see Fig.1, row 2). A left temporal and parieto-occipital dominant activation appears at lower SNRs then followed by a right temporal activation. The average Spearman’s correlation in the temporal and parieto-occipital areas in quiet is 0.06. In the theta EEG band (see Fig.1, rows 3 and 4), we observed a central activation, increasing with the SNR, reaching a maximum (averaged Spearman’s correlation around 0.04) between −0.5 and 2.5 dB SNR, then decreasing in quiet. Interestingly, an occipital-right dominance appears using the alpha EEG band at −6.5 db SNR, then followed by a centro-frontal dominance for SNR ranging between −5.5 and −1 dB SNR, finally reaching a maximum at −0.5 dB SNR with a fronto-left occipital activation. In the beta and low-gamma EEG bands, the cortical speech tracking at each electrode location was below the significance level (around 0.01). Note the decrease of the cortical speech tracking with EEG band frequency. Based on the spatial distribution of the cortical tracking of each speech representation, we defined a region of interest (ROI) for the delta (see Fig. 2, in red) and theta (see Fig. 2, in blue) frequency bands. A similar spatial distribution could be observed with the other models (see Fig Ap1 to Ap4 in the Appendix).

**Fig. 1.**
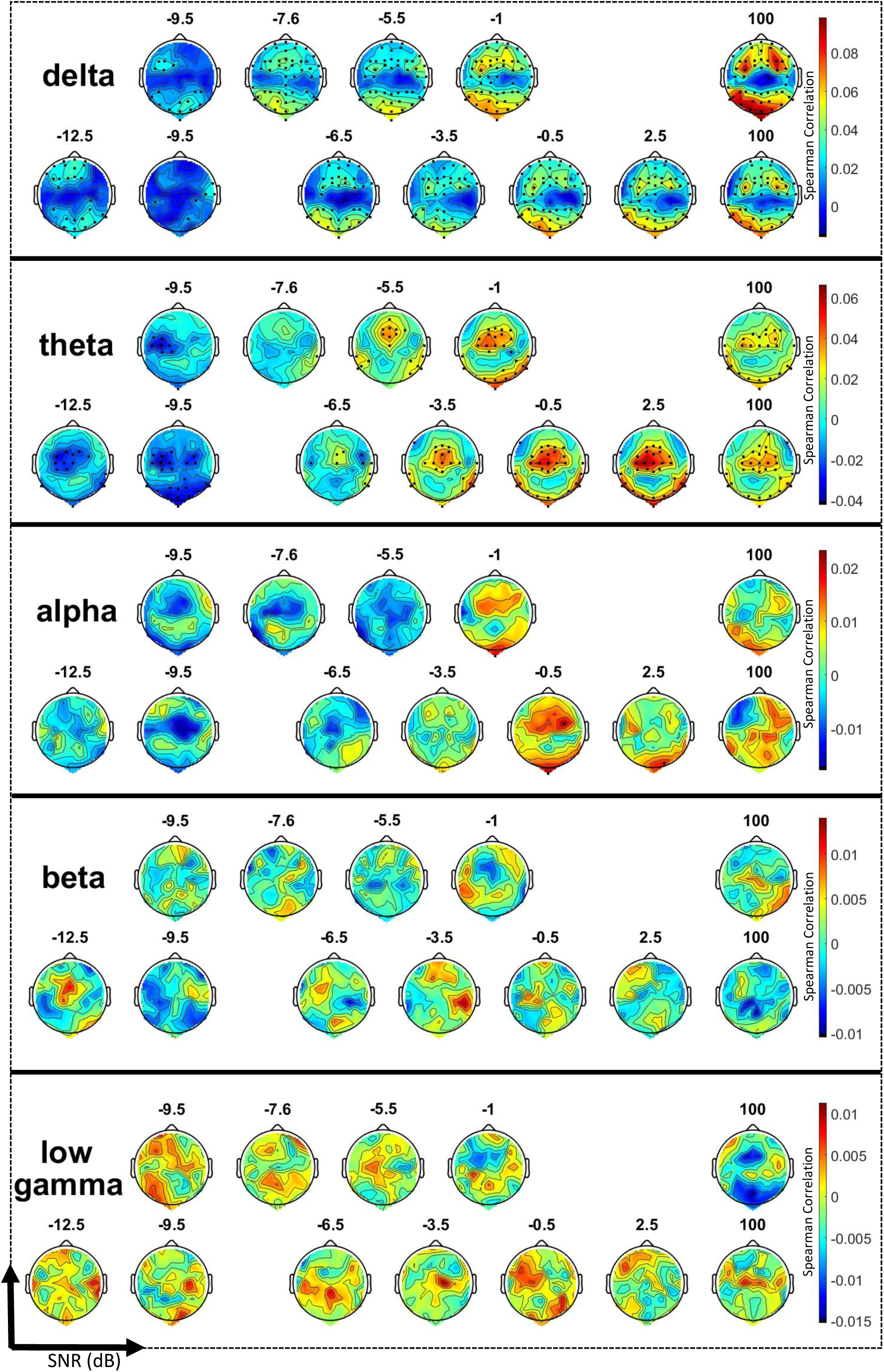
Spatial distribution of the cortical speech tracking at different levels of SNR over the scalp (64 electrodes) using a FS-model and either the delta (1-4 Hz), theta (4-8 Hz), alpha (8-15 Hz), beta (15-30 Hz) or low-gamma (30-45 Hz) EEG frequency band. For each frequency band, the first row represents the topographic map averaged over the participants of the first group (SNRs between −9.5 dB SNR and quiet); the second row represents the EEG prediction for the second group (SNRs between −12.5 dB SNR and quiet) using the same EEG frequency band. Channels with prediction above the level of significance (permutation test; p = 0.01) are indicated by a black dot. Note the increase of the speech cortical tracking with an increase in SNR in the lateralized fronto-central and occipital areas for the delta EEG band and in the central area for the theta EEG band, while almost none of the channels showed EEG predictions above the level of significance using the alpha, beta or low-gamma EEG bands.

**Fig. 2.**
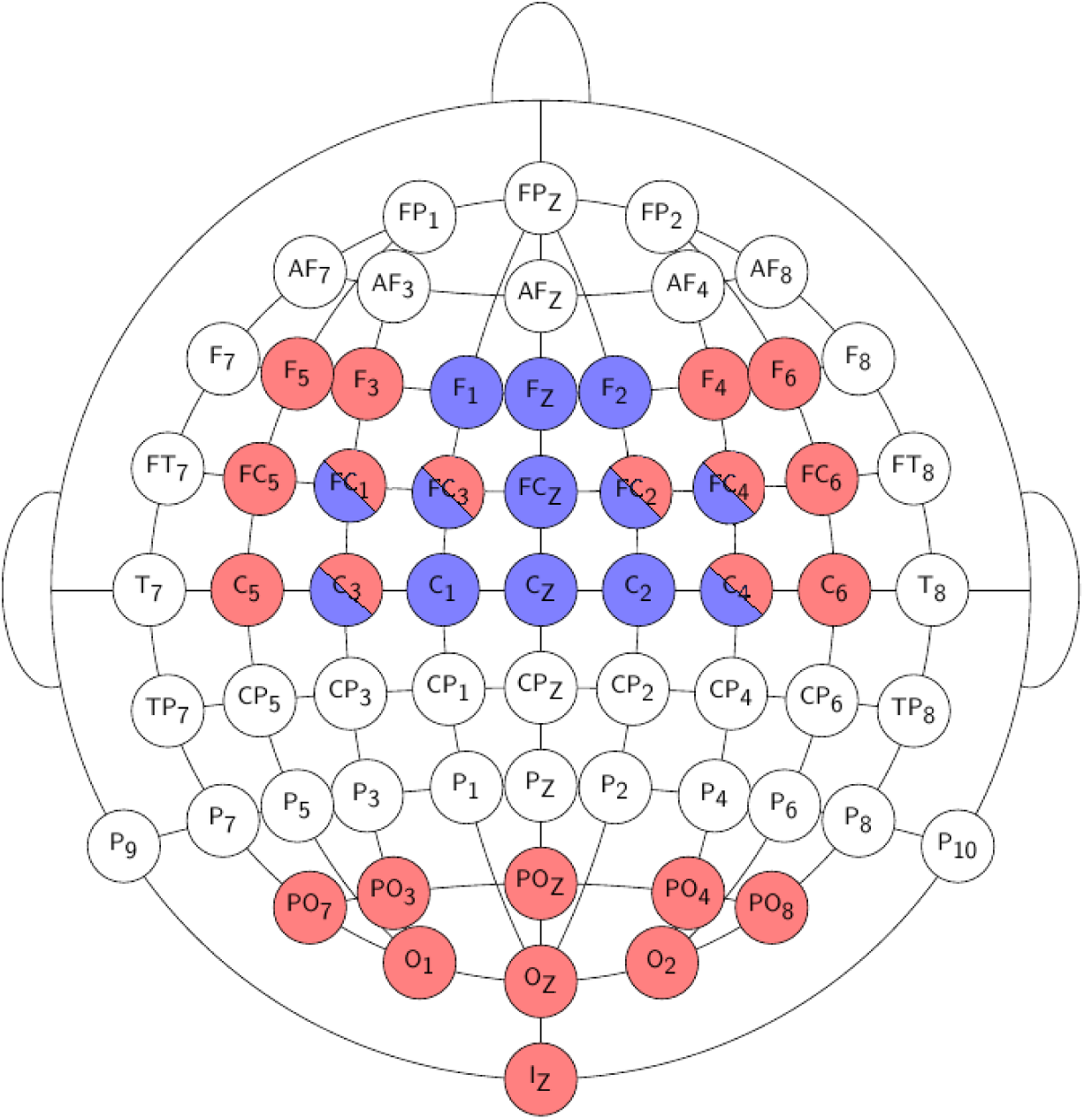
Electrode selection for the delta (in red) and theta (in blue) frequency bands. Note the fronto-temporal and occipital dominance of the delta band over the scalp while the theta activation is centralized in the frontal area.

### Group-level and individual cortical tracking in function of stimulus SNR

Both for group 1 and for group 2, the averaged correlation over the delta ROI increased with the SNR using each speech representation (see Fig. 3, rows 1 and 2). The averaged correlations for SNRs below −3.5 dB SNR (resp. below −6.5 dB SNR) were at the significance level using either the Env or Ph (resp. Fea) models. The Sgram and FS models showed correlations above the significance level for all SNRs. The FS model showed similar correlations at lowest SNRs as the Sgram model, but with a higher slope for both group 1 and group 2. In the delta band, the correlation in quiet increased with model complexity/feature level (Env → Sgram →… → FS), the highest correlation being reached using the FS model. Models based on low-level features (Env, Sgram and FS) showed a S-shape correlation over SNRs in the theta band (i.e., flat, followed by an increase, followed by a drop in quiet; see Fig. 3, rows 3 and 4), while the higher-level feature-based models presented a strictly monotonic increase in correlation over the SNR. The Ph and Fea models showed correlations above significance-level for SNRs above respectively −5.5 and −7.6 dB SNR for group 1 and above respectively −3.5 and −6.5 dB SNR for group 2. Interestingly, the Sgram and FS models showed a binary correlation trend, switching from significant negative correlation for the lowest SNRs (i.e., ≤ −9.5 dB SNR) to significant positive correlation for SNR above −6.5 dB SNR.

**Fig. 3.**
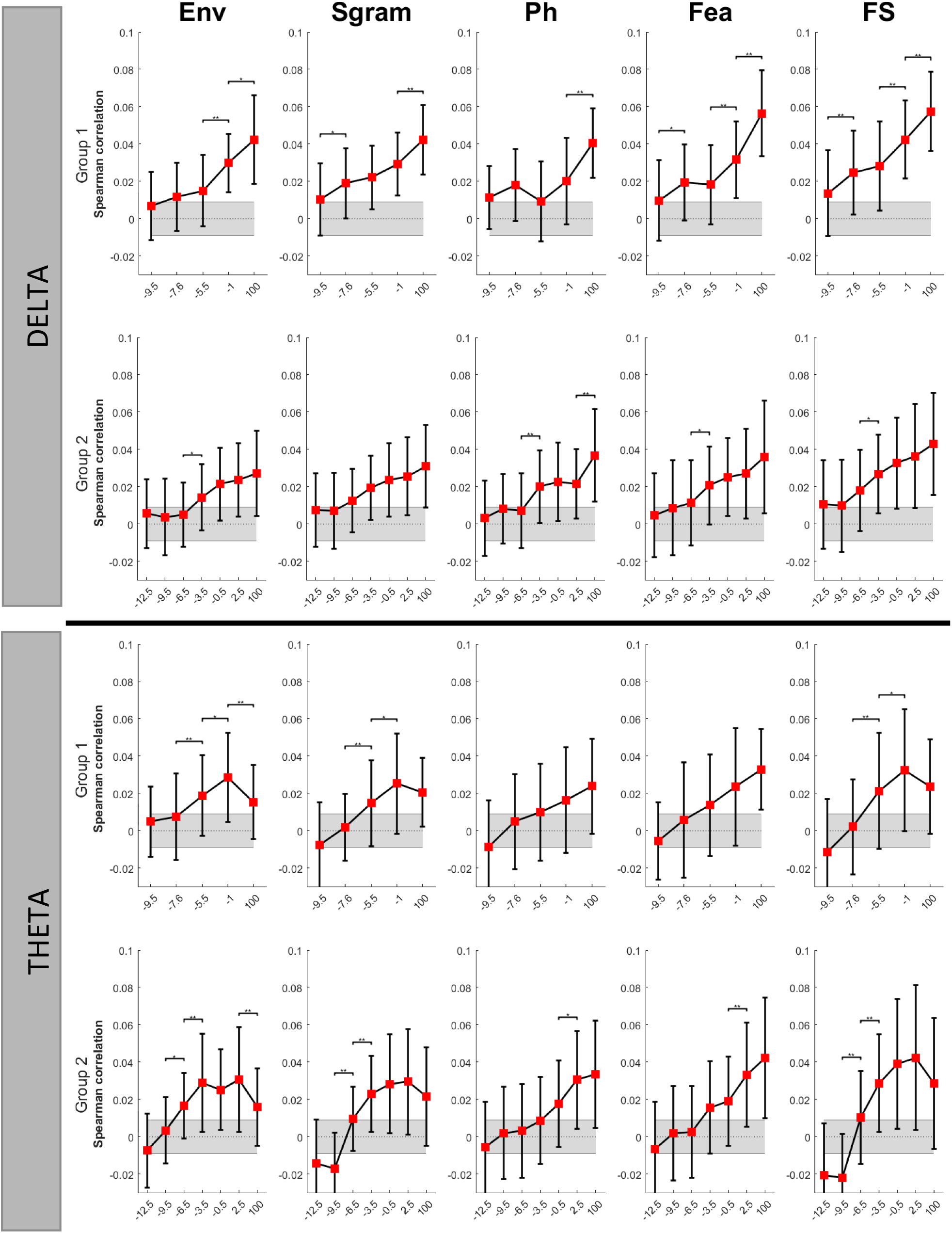
Averaged correlation over groups, frequency bands (rows) and speech representations (column). Note the monotonic increase in correlation for all the speech representations in each group in the delta band, and in the theta band for the Ph and Fea models. In the theta band, the models based on low-level speech features (Env, Sgram and FS) show a S-shape. In grey, the significance level (p = 0.01) calculated using a permutation test (1000 permutations) is shown.

In the appendix (see Fig. Ap6 and Ap7), we present the cortical speech tracking in function of SNR for each individual subject separately, using the delta-Env and delta-FS model or the theta-FS model respectively. At single-subject level, in the theta band, we observed an increase in cortical speech tracking with SNR (see Fig. Ap7), reaching a maximum around 0 dB SNR, then decreasing again. Sixteen out of the 19 participants showed a negative correlation between the actual and predicted EEG for lower SNRs. The averaged SNR at which the cortical speech tracking switched from a negative to a positive value is −7.2 ± 1.8 dB SNR (hereafter named FS-zeroCrossing).

### Predicting the SRT using the cortical tracking over SNRs

We finally evaluated to what extent we could predict the SRT using each model. We calculated (1) the number of participants for whom a CT could not be calculated from the EEG (see NA in Fig. 4) and (2) the percentage of participants for whom the absolute difference between the SRT and the CT was below 1dB SNR and 2dB SNR respectively (see Table 1).

**Fig. 4.**
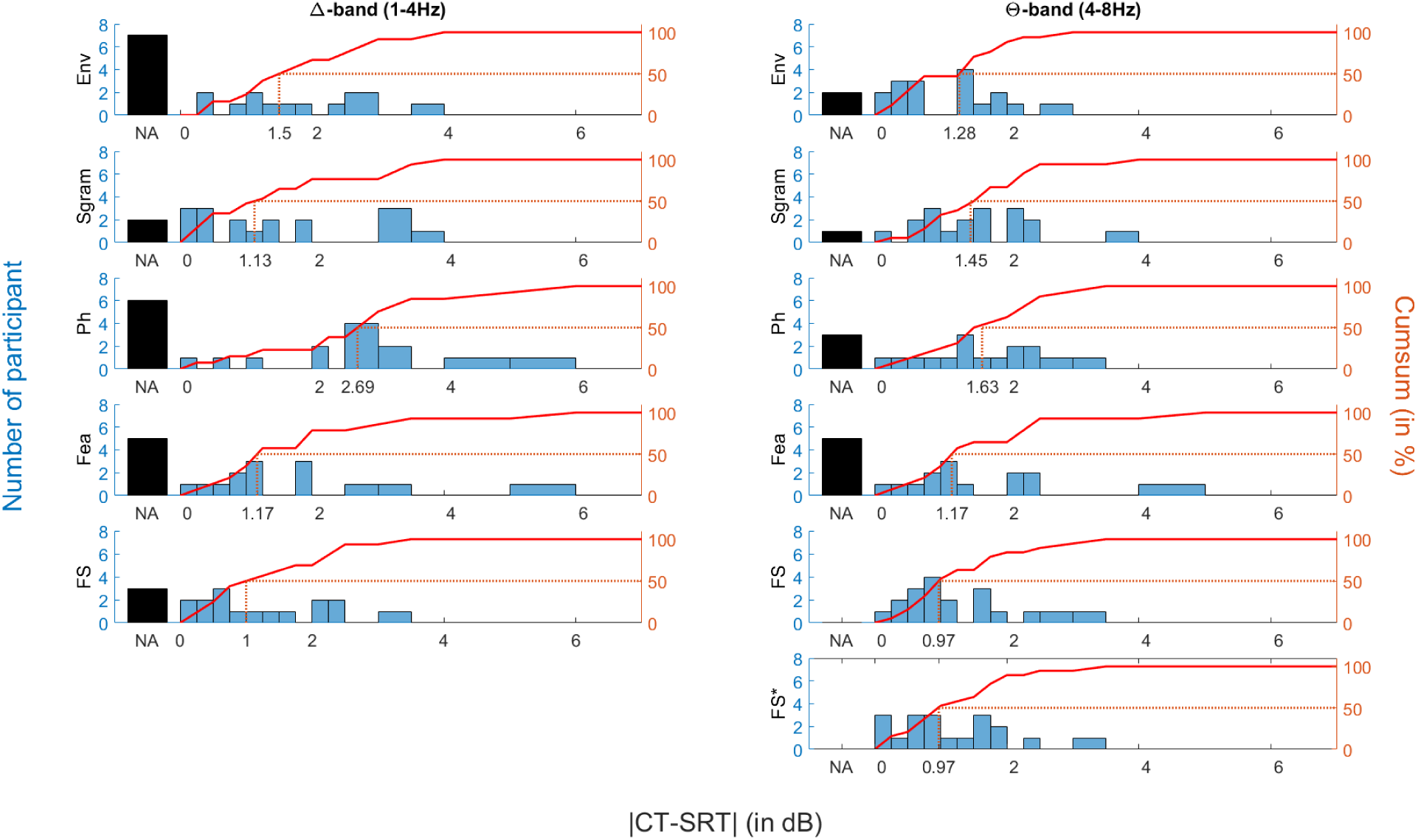
Histogram of the difference between the CT derived from the EEG responses to speech and the behavioral SRT, in function of speech representation (rows) and frequency band (columns). *FS\** stands for FS-zeroCrossing. The left y-axis shows the number of participants (in blue); the right y-axis shows the corresponding cumulative percentage of participants (in red). The dashed line represents the relative difference at which we can predict at least 50% of the participants’ SRTs. The black bar (NA) shows the number of participants for which the CT could not be calculated. Interestingly, the models based on the delta EEG band yielded a higher NA score. As expected, using the theta EEG band, models presenting a S-shape in their speech cortical tracking over SNR (i.e., Env, Sgram and FS) yielded both a lower NA score and a smaller difference between the CT and SRT.

**Table 1.**
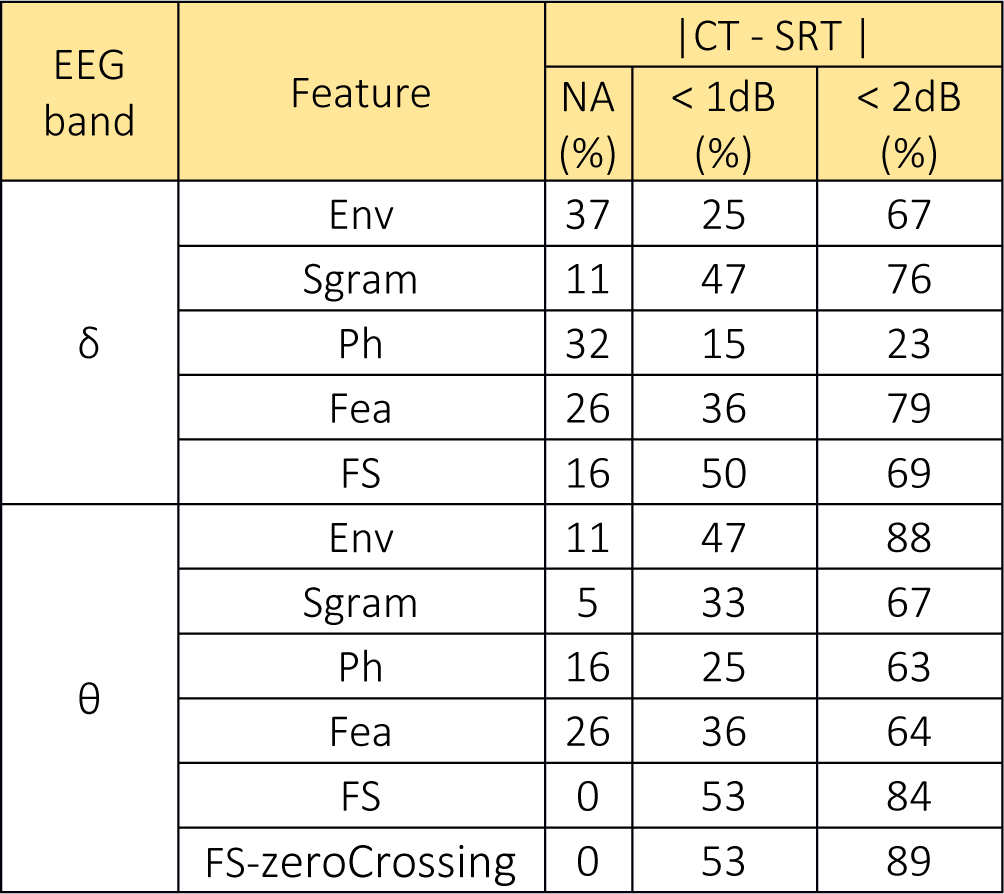
The percentage of participants for whom the CT score could not be measured is shown in the column titled NA. For the participants for whom the CT score could be measured, we computed the number of participants presenting an absolute difference (in dB) between the objective CT and the behavioral SRT of less than 1dB (see the fourth column) and 2dB (see the fifth column). Note that only 16% of the participants (i.e., 25% of the 63%) present an absolute difference of less than 1 dB with the Env-delta model, while 53% for the FS-theta model.

| EEG band | Feature | \|CT - SRT\| |  |  |
| --- | --- | --- | --- | --- |
|  |  | NA (%) | < 1dB (%) | < 2dB (%) |
| $\delta$ | Env | 37 | 25 | 67 |
|  | Sgram | 11 | 47 | 76 |
|  | Ph | 32 | 15 | 23 |
|  | Fea | 26 | 36 | 79 |
|  | FS | 16 | 50 | 69 |
| $\theta$ | Env | 11 | 47 | 88 |
|  | Sgram | 5 | 33 | 67 |
|  | Ph | 16 | 25 | 63 |
|  | Fea | 26 | 36 | 64 |
|  | FS | 0 | 53 | 84 |
|  | FS-zeroCrossing | 0 | 53 | 89 |

The delta-Env model illustrated the highest fraction (37%) of participants for whom a CT score could not be calculated. The theta models showed the lowest number of participants whom a CT score could not be calculated. In particular, a CT score could be calculated for each participant using the theta-FS model.

With the delta Env-model, 3/19 participants illustrated a difference between CT and SRT scores of less than 1 dB, while 8/19 participants showed a difference of less than 2dB (see Fig. 4, row 1, first column). Using the theta-FS model, 10/19 participants had a difference of less than 1 dB, and 16/19 participants presented a difference of less than 2 dB (see Fig. 4, row 4, second column). Interestingly, using the FS-zeroCrossing model, 11/19 participants presented a difference between the CT and the SRT of less than 1dB, and 18/19 with a difference of less than 2dB (see Fig. 4, last row).

## DISCUSSION

We showed that the combination of both low- and higher-level speech features within a model could improve the prediction of speech intelligibility from the EEG, compared to just the standard speech envelope. For the delta band, the FS-model, which combines the spectrogram and phoneme representations, yielded higher correlations between actual and predicted EEG in the no-noise condition than the Env-model, while the correlation at the lowest SNRs was the same, suggesting a better sensitivity of this model to the level of speech understanding (i.e., a higher slope of the cortical speech tracking over SNR). Moreover, when comparing the behavioural SRT with its objective counterpart (the CT), for the theta FS-model, the difference between SRT and CT was under 2 dB for more than 80% of the participants. Part of the remaining difference can be explained by the test-retest difference of the behavioral measure, which is around 2 dB (Francart et al. 2011; Decruy, Das, et al. 2018). We could hypothesize that the FS model benefits from the integration of both low-level and higher-level speech features so the delta-FS model benefits from the good monotonicity of the delta-Fea model (with a small contribution of the delta-Sgram model) and the theta-FS model can better predict the SRT thanks to its S-shape that probably derived from the theta-Sgram model. Future research should evaluate the inclusion of higher-level speech features such as words or semantic information, phoneme-level surprisal, entropy, or phoneme onset in the FS-model (Broderick et al. 2018; Brodbeck et al. 2018).

The spatial distribution of the EEG predictability over the scalp showed a lateralized fronto-temporal and occipital dominance for the delta EEG band, and a central centro-frontal dominance for the theta EEG band. Recently, Destoky et al. (2019) evaluated cortical tracking of the envelope using both EEG and MEG and showed a group-level coherence map between the brain responses and speech envelope with a similar pattern. A lateralized fronto-temporal brain coherence has also been observed in the delta EEG band by Mukherjee et al. (2019). Studying the change in activation with stimulus SNR in these areas of interest, we showed that in the delta band each model yielded a monotonic increase of the cortical speech tracking with SNR. In the theta band this could also be observed using the Ph or Fea model, but the models based on low-level speech features (Env and Sgram) showed an S-shape with a maximum correlation around 0dB SNR. We hypothesize that these two frequency bands track different features of the speech signal in the brain, as suggested by Ding & Simon (2014). The drop in correlation between a condition with limited noise (0 dB SNR) and no noise might be due to a drop in listening effort or attention (Das et al. 2018). Future studies should be conducted to better understand and characterize this difference.

As suggested by Vanthornhout et al. (2018) using a backward model to predict speech intelligibility using the speech envelope, we here fitted a sigmoid to the cortical speech tracking as s function of SNR in order to predict the CT. It is however important to stress that fitting a sigmoid is not an easy step (i.e., it is difficult to fit a four-parameter-curve using a limited set of measures). Indeed, it led to a number of participants for whom the CT score could not be measured (see the NA score in Table 1, around 17% in average over the different models; note that in Vanthornhout et al. (2018), this NA proportion is at 21%). A way to mitigate this risk would be to increase the number of repetitions for each SNR (i.e., 4 repetitions for group 1 and 3 for the group 2 in the present study) at the cost of session time. To overcome this issue, we here suggested a FS-zeroCrossing score that provides similar results as the FS-theta model (i.e., 53% and 89% of the participants with an absolute difference between the CT and SRT of less than 1dB and 2dB respectively) without the cost of a sigmoid fitting. One possible intuitive explanation of why this FS-zeroCrossing seems to correspond to the behavioral SRT is that, at the SRT, half of the EEG is correlated with the speech (i.e., the brain tracks the speech features of the 50% words that can be understood); half is uncorrelated, counterbalancing the correlated brain activity. Above that threshold, the proportion of correlated vs uncorrelated brain signals is higher than one, so the resulting correlation is above the level of chance.

The paradigm and subject population was well-controlled in this study. However, for clinical application of our measure, a number of potential factors should be considered. Some factors are related to the participant (i.e., how does change in the participant’s state over time impact our measure?) or from group-specific characteristics (i.e., does this measure work for the whole population?). Indeed, little is known about the impact of factors such as attention or motivation on the cortical tracking of speech features. However, Kong et al. (2014) found that while cortical envelope tracking in a single-talker paradigm can be affected by task and attention, it still increases with stimulus SNR (Vanthornhout et al. 2018). Moreover, we here evaluated our measure on a specific cohort, encompassing young university students without any known pathologies. Education may affect comprehension. In addition, recent studies have shown that cortical tracking of speech can increase with age (Presacco et al. 2016; Decruy et al. 2018), but still consistently increases with stimulus SNR. Future studies should evaluate the impact of these and other factors (Blank & Davis 2016; Di Liberto et al. 2018) on our measure of speech intelligibility since a large proportion of hearing-impaired population is elderly and has a varied educational background.

While we here propose a novel clinical test that works under well-defined circumstances and stimuli, it is important to keep in mind that the measured level of cortical tracking is a proxy for intelligibility, rather than a direct measurement. We do not make any claims regarding causality. A number of factors could explain the sensitivity of our metric to the level of speech understanding. For example, Peelle and Davis (2012) indicated that intelligible speech engages a broader network than non-speech/unintelligible sounds. Consequently, one possibility is that stronger EEG predictions would arise for speech stimuli that are more intelligible, simply because there is “more brain activity” correlated with the stimulus. Another possibility is that the same brain activity becomes more “precise” or synchronized with the stimulus. In addition, in our study, we predicted the EEG responses to speech in different noisy conditions (Matrix sentences) using the TRFs trained with speech in silence (Story). One possible caveat is that the EEG prediction results reflect the similarity (in other words, the matching or consistency) of the EEG responses to a model of clear speech perception, rather than a difference in the strength of the encoding of the underlying responses. We tested this hypothesis by attempting the reverse: training the different models using each group’s lowest SNRs Matrix data (i.e., by concatenating the different 2-min presentations of the lowest SNR) and then testing it on the remaining SNR conditions. If the model in this case encodes the similarity with speech in noise, we hypothesize that high EEG prediction results would be observed in the lowest SNRs, then decreasing with increasing SNR. However, no cortical tracking to the speech could be observed in this scenario (see Fig. Ap8) suggesting that the EEG predictions reflect the intensity of the speech responses and not similarities between noisy conditions.

Ding and Simon (2013) showed that the TRF depends on the SNR, so a model trained on clean speech could be inappropriate to predict speech in noise. We investigated this by training the model on noisy speech instead of speech in silence (Fig. Ap9 and Fig. Ap10). While we agree that the level of noise can induce changes in the TRF, the speech cortical tracking could be extracted in all training conditions and resulted in similar topographies as in our original experiment. Considering the application of our method in a clinical setting, we decided to train our model using speech in silence data. First, this allows to use patient’s brain response to a verbal instruction in order to train a model prior the testing part, and to establish baseline neural tracking for the patient. Second, this could increase participants’ motivation by presenting a fully understandable fairy tale (in our study, the story “Milan”) rather than a sequence of sentences. However, further research is needed to establish the most efficient clinical protocol.

Integrating low and higher-level speech features, the novel measures proposed in this study provide an objective measure of speech intelligibility and open the doors to automatic hearing aid and cochlear implant (Somers et al. 2019) parameter adaptation relying on auditory brain responses and requiring minimum intervention of a clinical expert. Future research should evaluate the potential of this new measure in an extended cohort in clinical conditions and study the impact of parameters like the semantic context, participant’s age, hearing loss, test-duration and number of electrodes included in the recording.

## ACKNOWLEDGEMENT

The authors would like to thank Prof. Hugo Van hamme for providing the phoneme segmentation. Financial support was provided by the KU Leuven Special Research Fund under grant OT/14/119 to Tom Francart. This project has received funding from the European Research Council (ERC) under the European Union’s Horizon 2020 research and innovation program (grant agreement No 637424, ERC starting Grant to Tom Francart). Research funded by a PhD grant of the Research Foundation Flanders (FWO) for Jonas Vanthornhout (1S10416N) and Eline Verschueren (1S86118N). The authors declare no conflict of interest

