## Supplementary Material for "Predicting individual speech intelligibility from the cortical tracking of acoustic- and phonetic-level speech representations"

### APPENDIX

Spatial distribution of the cortical tracking of different EEG bands

The following figures show the same results as figure 1 in the main text, but now for different models: Env, Sgram, Ph and Fea.

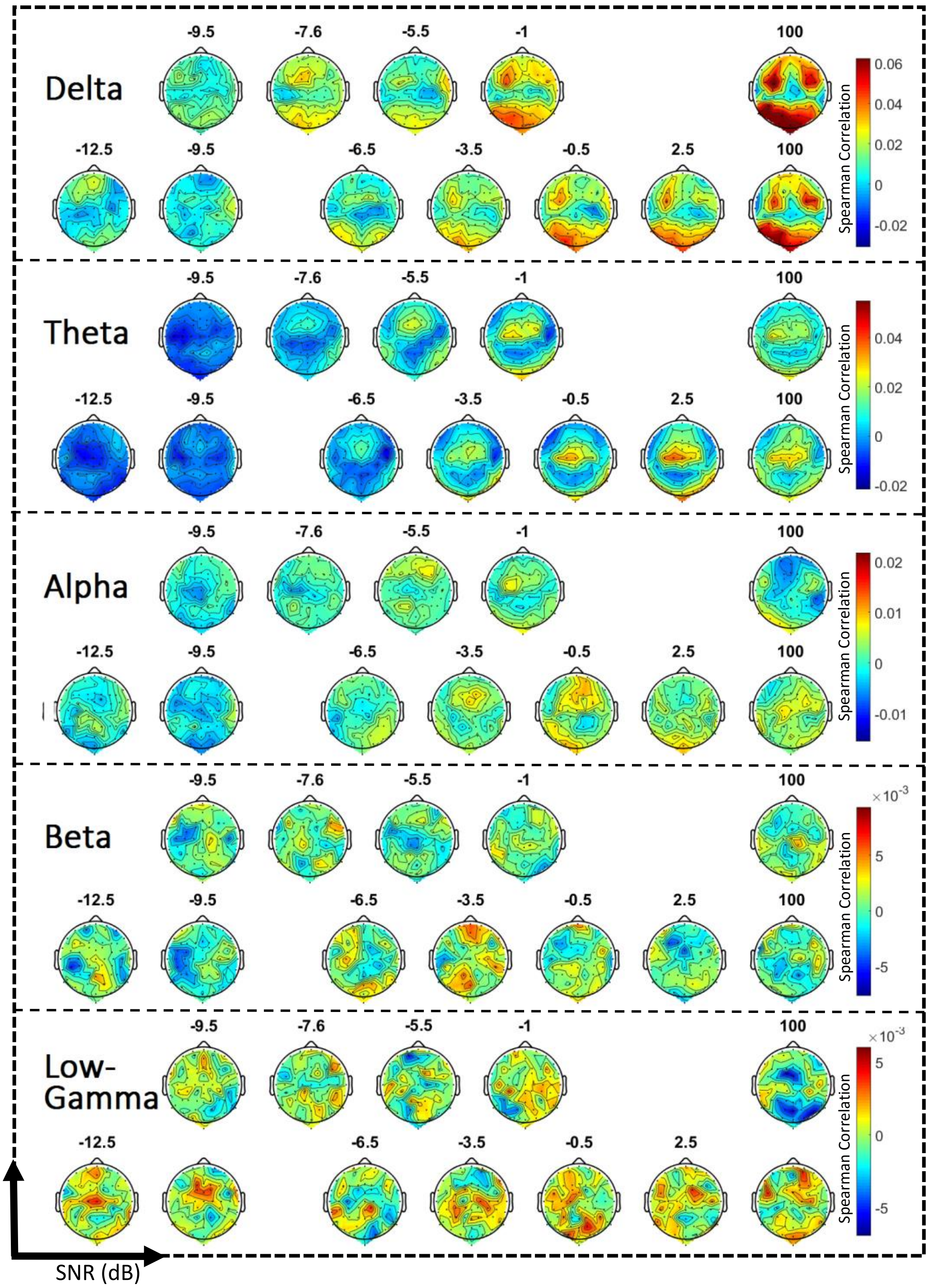


Fig. Ap1 - Spatial distribution of cortical speech tracking at different levels of SNR over the scalp (64 electrodes) using an **Env-model** and either the delta (1-4 Hz), theta (4-8 Hz), alpha (8-15 Hz), beta (15-30 Hz) or low-gamma (30-45 Hz) EEG frequency band. For each frequency band, the first row represents the topographic map averaged over the participants of the first group (SNRs between -9.5 dB SNR and quiet); the second row represents the EEG prediction for the second group (SNRs between -12.5 dB SNR and quiet) using the same EEG frequency band.

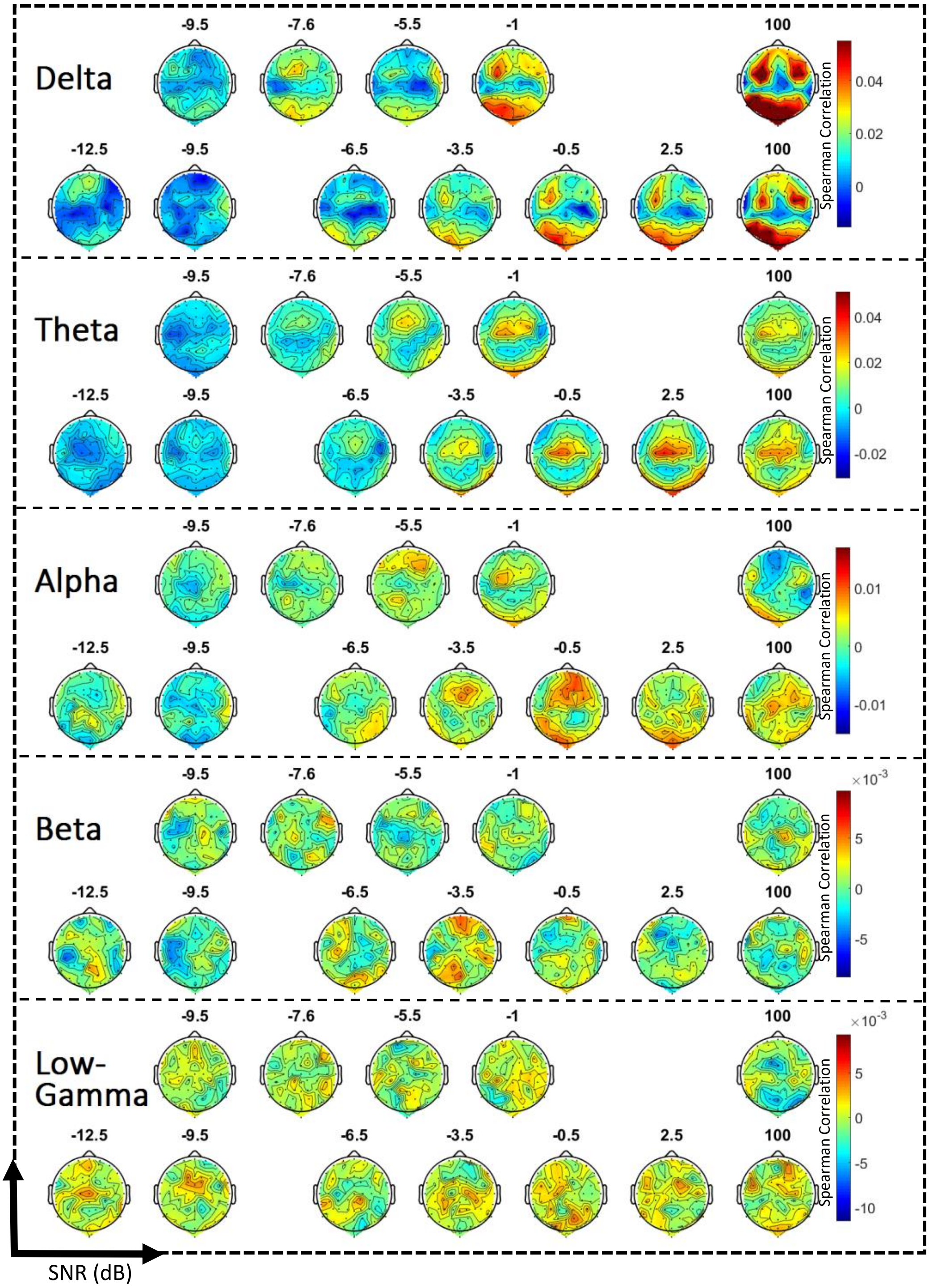


Fig. Ap2 - Spatial distribution of the cortical speech tracking at different levels of SNR over the scalp (64 electrodes) using a **Sgram-model** and either the delta (1-4 Hz), theta (4-8 Hz), alpha (8-15 Hz), beta (15-30 Hz) or low-gamma (30-45 Hz) EEG frequency band. For each frequency band, the first row represents the topographic map averaged over the participants of the first group (SNRs between -9.5 dB SNR and quiet); the second row represents the EEG prediction for the second group (SNRs between -12.5 dB SNR and quiet) using the same EEG frequency band.

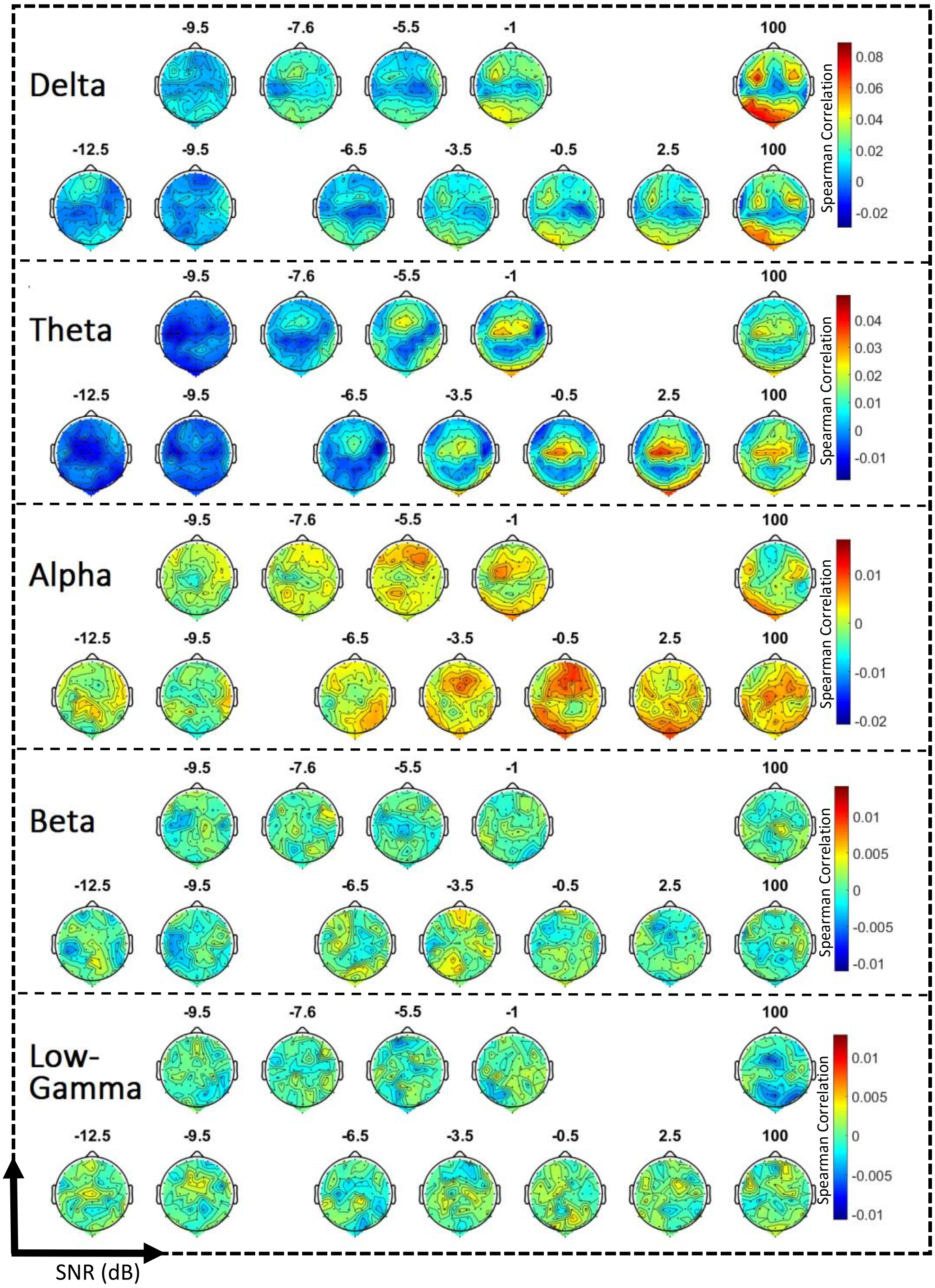


Fig. Ap3 - Spatial distribution of the cortical speech tracking at different levels of SNR over the scalp (64 electrodes) using a **Ph-model** and either the delta (1-4 Hz), theta (4-8 Hz), alpha (8-15 Hz), beta (15-30 Hz) or low-gamma (30-45 Hz) EEG frequency band. For each frequency band, the first row represents the topographic map averaged over the participants of the first group (SNRs between -9.5 dB SNR and quiet); the second row represents the EEG prediction for the second group (SNRs between -12.5 dB SNR and quiet) using the same EEG frequency band.

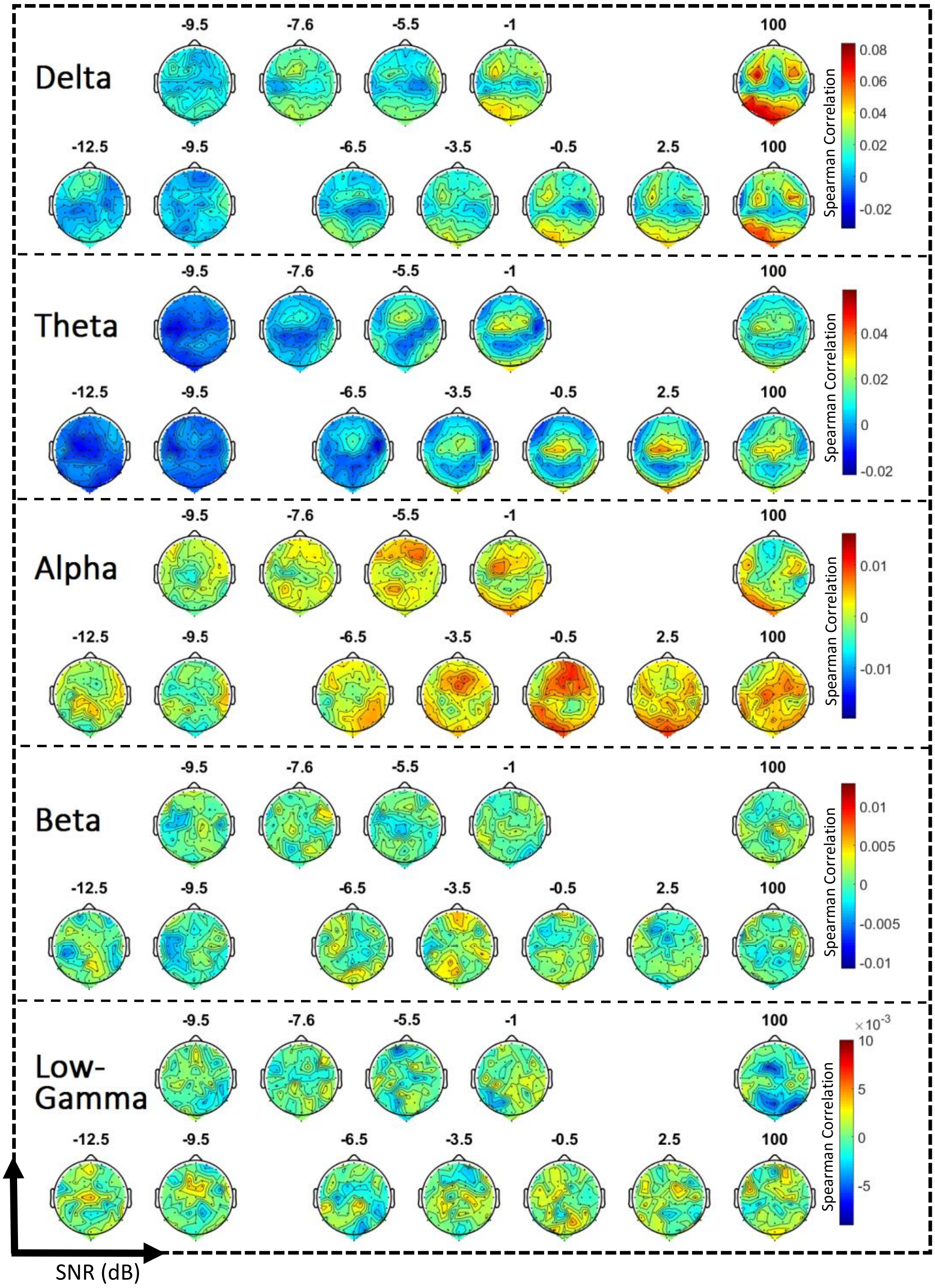


Fig. Ap4 - Spatial distribution of the cortical speech tracking at different levels of SNR over the scalp (64 electrodes) using a **Fea-model** and either the delta (1-4 Hz), theta (4-8 Hz), alpha (8-15 Hz), beta (15-30 Hz) or low-gamma (30-45 Hz) EEG frequency band. For each frequency band, the first row represents the topographic map averaged over the participants of the first group (SNRs between -9.5 dB SNR and quiet); the second row represents the EEG prediction for the second group (SNRs between -12.5 dB SNR and quiet) using the same EEG frequency band.

Monotonicity of the cortical tracking over SNRs

*Methods* - To compare each model's ability to track the participant's speech intelligibility, we assessed the extent to which cortical tracking increased monotonically with the stimulus SNR. We defined a monotonicity score (MS) as follows: for a stimulus $SNR_i$, we compared the associated Spearman's correlation $\rho(SNR_i)$ with ones obtained at lower and higher SNRs (i.e., $\rho(SNR_1)$ … $\rho(SNR_{i-1})$, $\rho(SNR_{i+1})$… $\rho(SNR_N)$) as well as with the significance level (see below). If $\rho(SNR_i)$ was above the significance level (see below), we incremented the monotonicity score as follows:

$$MS_r = MS_r + \frac{\left(\sum_{k=1}^{i-1} \exists\left(\rho(SNR_k) \leq \rho(SNR_i)\right) + \sum_{k=i+1}^{N} \exists\left(\rho(SNR_k) \geq \rho(SNR_i)\right)\right)}{N-1}$$

i.e., for $SNR_i$ with a Spearman correlation $\rho(SNR_i)$ above the significance level, the monotonicity score of the participant r (i.e., $MS_r$) was increased by the ratio of the number of Spearman's correlations of lower SNR below or equal to $\rho(SNR_i)$ added to the ratio of the number of Spearman's correlations of higher SNR above or equal to $\rho(SNR_i)$. If $\rho(SNR_i)$ was below the significance level, the score was not increased.

*Results* - The highest averaged monotonicity score (i.e., mean ± std, 73 ± 20%) was obtained in the delta EEG frequency band with the FS-model (see Fig. Ap5). This model outperformed the Env (64.5 ± 20 %; WSRT, $W(19) = 30.5$, $p < 0.05$, two-tailed test) and Sgram (67 ± 21 %; WSRT, $W(19) = 11$, $p < 0.01$, two-tailed test) models. The monotonicity score decreased with increasing EEG frequency bands. Comparing the FS and Env models at single-subject level, we showed that the FS model outperformed the Env-model in 14/19 participants, and two of the remaining showing no difference in monotonicity score between the two models (see Fig. Ap6).

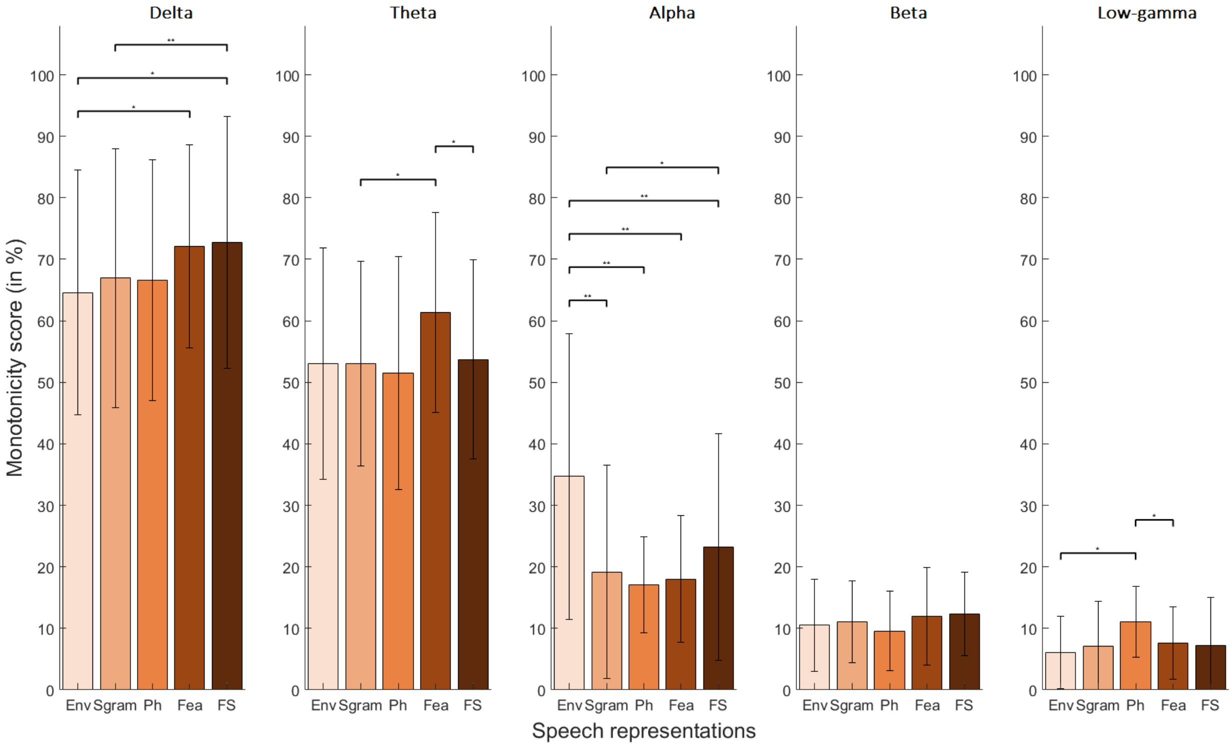


Fig. Ap5 – Averaged monotonicity scores for the different models and EEG frequency bands. This score quantifies to what extent a model provides a monotonic increase of its cortical speech tracking with SNR for both groups. Note the decrease of this score with the frequency band. The highest score is reached for the FS-model in the delta band.

Individual cortical speech tracking in function of SNR

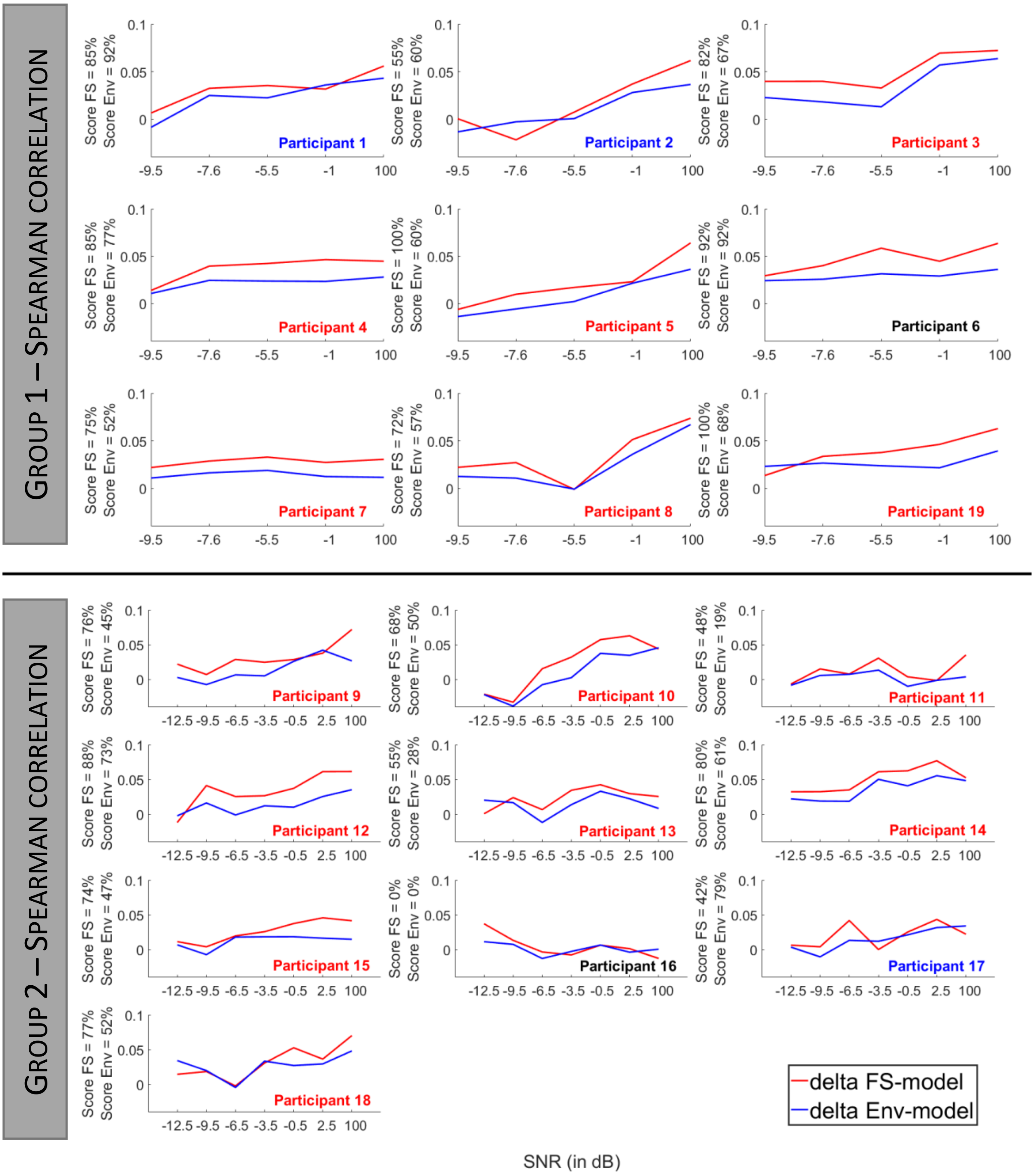


Fig. Ap6 – Individual cortical speech tracking over different SNRs in the delta band for the Env (in blue) and FS (in red) models. For each participant, the monotonicity score is shown on the y-axis. Participant numbers for whom the monotonicity score was higher for the Env (resp. FS) model are shown in blue (resp. in red).

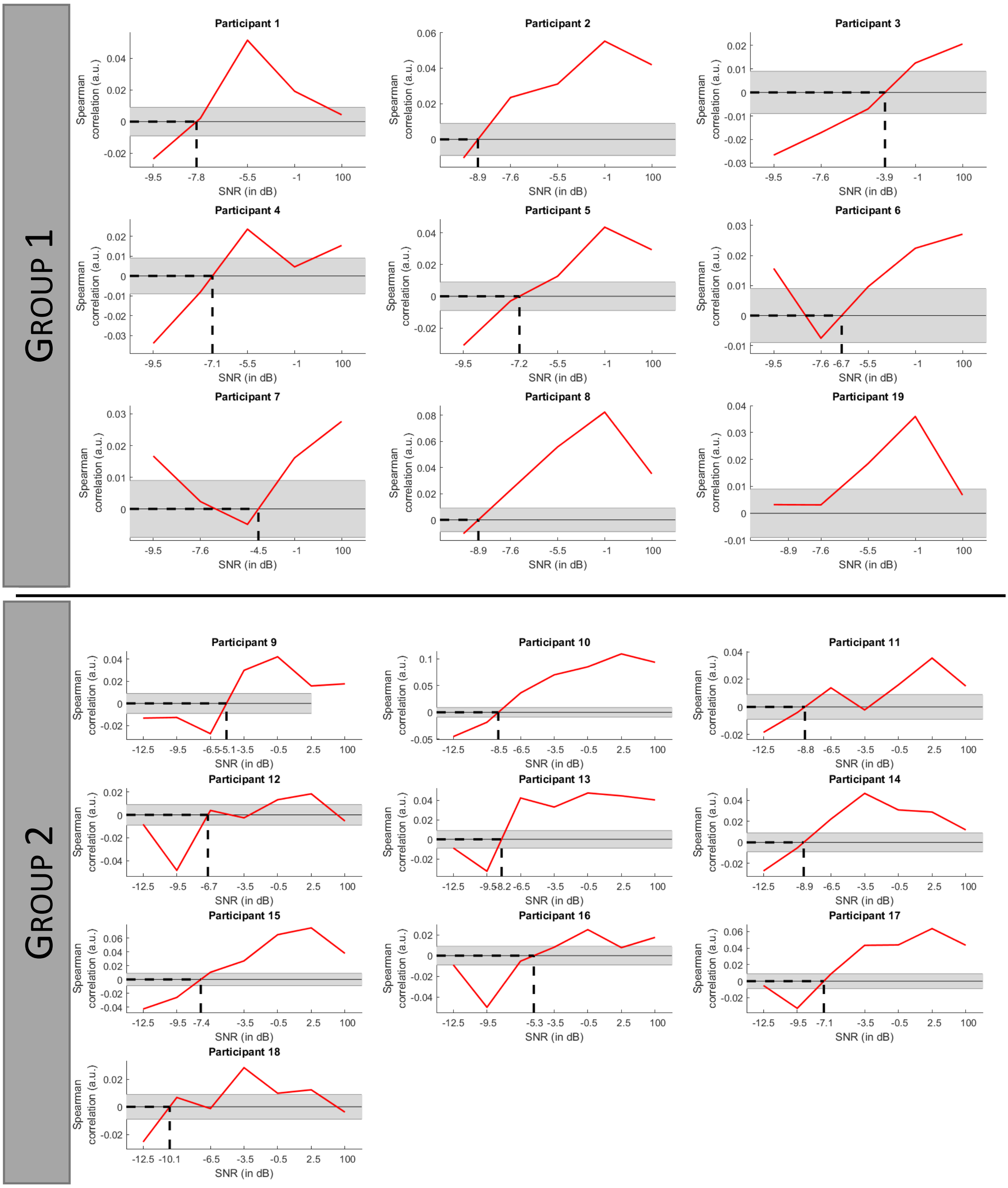


Fig. Ap7 – Individual cortical speech tracking over different SNRs in the theta band for the FS model. For each participant, the SNR at which the Spearman correlation switches from negative to positive is shown with a dashed line.

### Investigation of the effect of the set of data used to train the model on topographies

In the paper, the models are trained using clean speech (the Story condition) and evaluated on the Matrix in noise condition. As Ding & Simon (2013) showed that the TRF depends on the SNR, we investigated the effect of making other selections for the training/testing data.

We repeated the analysis using three different scenarios:

- Analysis A (Fig Ap8): we built a grand-average decoder using all the presentations at the lowest SNR (e.g., the 3 presentations of Matrix at -12.5 dB SNR) and then tested it at higher SNRs.
- Analysis B (Fig Ap9): we built a grand-average decoder using only one presentation of all the SNRs (e.g., one presentation of Matrix at -12.5, -9.5, -7.6, -6.5, -0.5, 1, dB SNR and no-noise) and then tested this model on the remaining presentations.
- Analysis C (Fig Ap10): we built a grand-average decoder for each SNR, using one presentation of each SNR and then tested the model on the remaining presentations of the same SNR.

Analysis A serves to show that the EEG predictions results do not reflect the similarity (in other words, the matching or consistency) of the EEG responses to a model of clear speech perception, rather than a difference in the strength of the the underlying neural responses. Analyses B and C serve to show that we can measure the speech cortical tracking using a model trained on speech in noise data, taking into account that TRFs may be different for noisy speech.

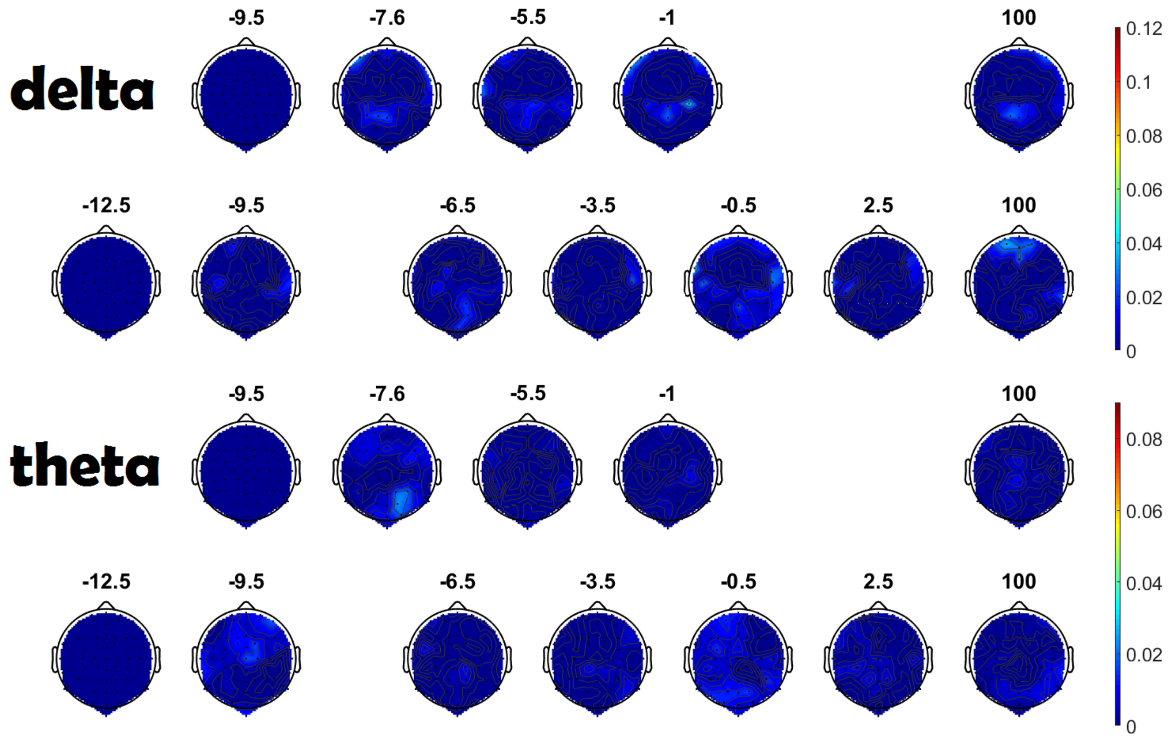


Fig. Ap8 – Topographies for a model trained using all the presentations at the lowest SNR (i.e., for group 1, the 4 presentations of Matrix sentences at -9.5 dB SNR; for group 2, the 3 presentations of Matrix sentences at -12.5 dB SNR). We used an FS-model and either the delta (1-4 Hz; first and second rows) or theta (4-8 Hz; third and fourth rows) EEG frequency band. For each frequency band, the first row represents the topographic map averaged over the participants of the first group (SNRs between -9.5 dB SNR and quiet); the second row represents the EEG prediction for the second group (SNRs between -12.5 dB SNR and quiet) using the same EEG frequency band.

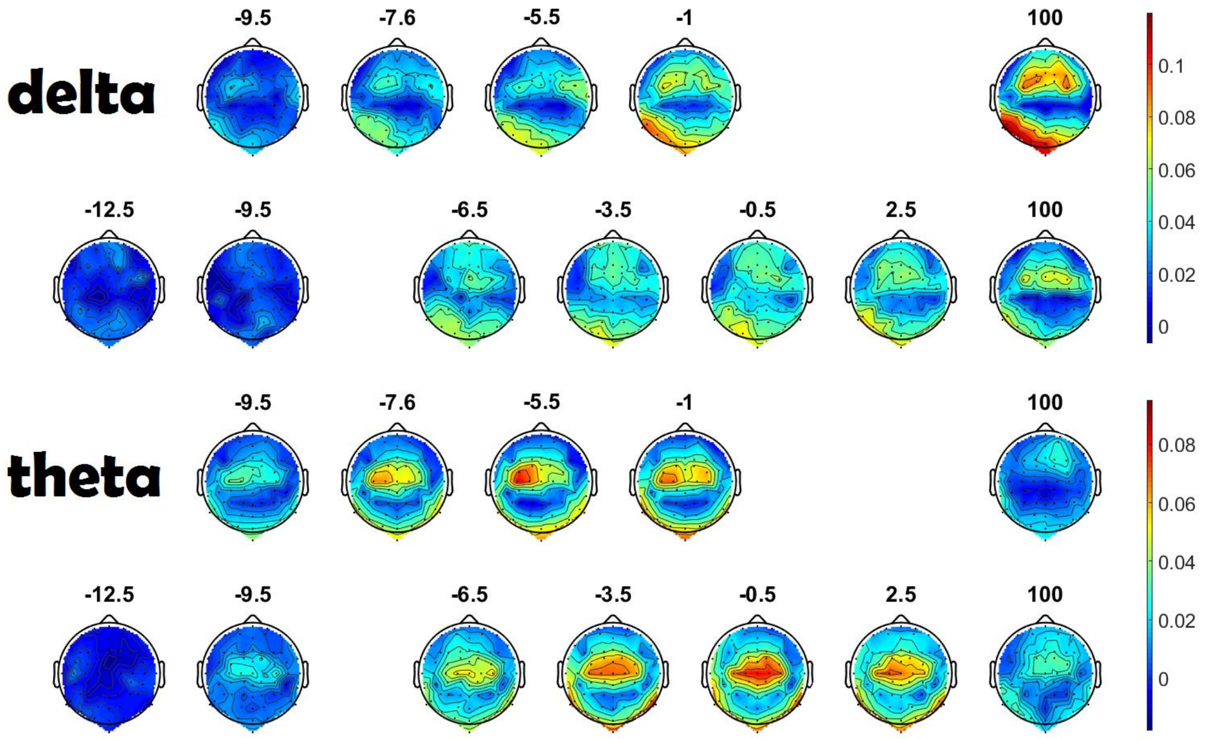


Fig. Ap9 – Topographies for a model trained on Matrix in noise data, using one model for all SNRs. The model was trained using one presentation of all the different SNRs (e.g., for group 2, one 2-min presentation of Matrix sentences at -12.5 dB SNR, one 2-min presentation of Matrix sentences at -9.5 dB SNR, one 2-min presentation of Matrix sentences at -7.6 dB SNR, … and one 2-min presentation of Matrix sentences in the no-noise condition). The model was then tested on the remaining presentations. We used an FS-model and either the delta (1-4 Hz; first and second rows) or theta (4-8 Hz; third and fourth rows) EEG frequency band. For each frequency band, the first row represents the topographic map averaged over the participants of the first group (SNRs between -9.5 dB SNR and quiet); the second row represents the EEG prediction for the second group (SNRs between -12.5 dB SNR and quiet) using the same EEG frequency band.

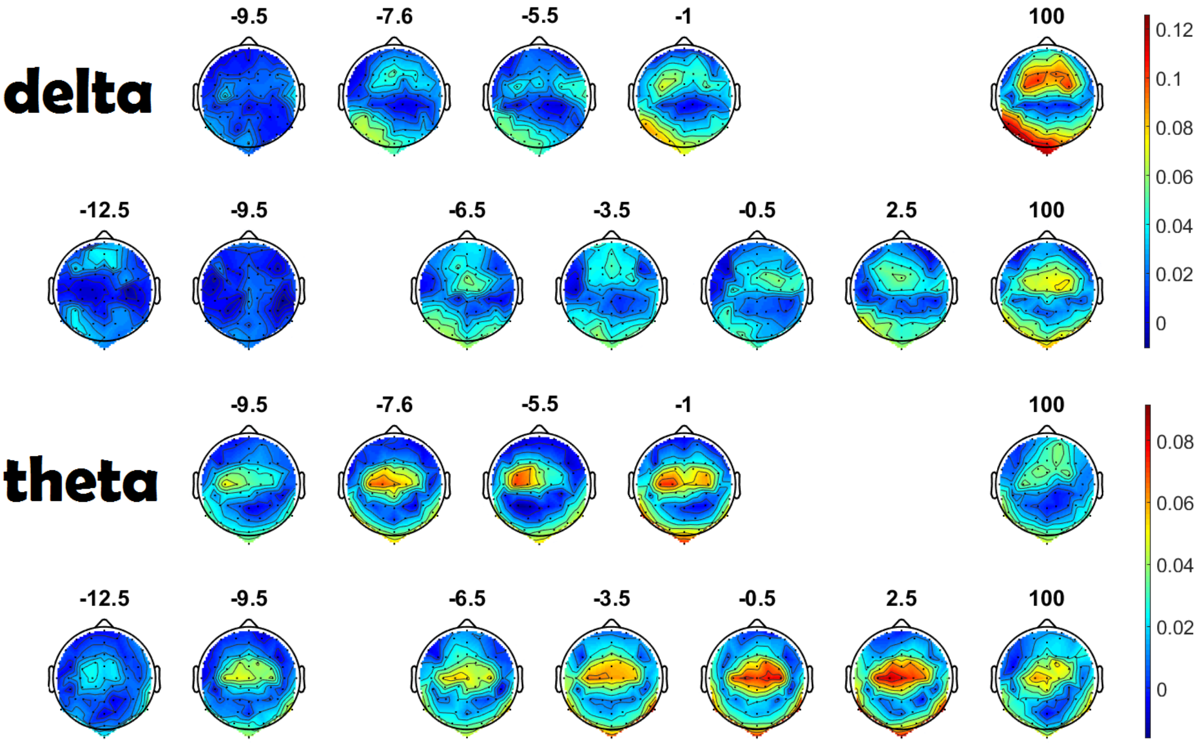


Fig. Ap10 - Topographies for individual models trained trained on Matrix in noise data, using a separate model for each SNR. Each model was trained using one presentation at its SNR (e.g., one 2-min presentation of Matrix sentences at -9.5 dB SNR) and then tested on the remaining presentations of the same SNR. We used a FS-model and either the delta (1-4 Hz; first and second rows) or theta (4-8 Hz; third and fourth rows) EEG frequency band. For each frequency band, the first row represents the topographic map averaged over the participants of the first group (SNRs between -9.5 dB SNR and quiet); the second row represents the EEG prediction for the second group (SNRs between -12.5 dB SNR and quiet) using the same EEG frequency band.

### Impulse and step responses of the pipeline

Figure Ap11 shows the impulse and step responses of the entire pipeline used for processing the EEG data in this study. See de Cheveigné and Nelken, Neuron, 2019, for in-depth clarifications on the risks of filtering. In our study, we could hypothesize that the impact of filter artefacts are minimal because we are using EEG prediction and correlation values.

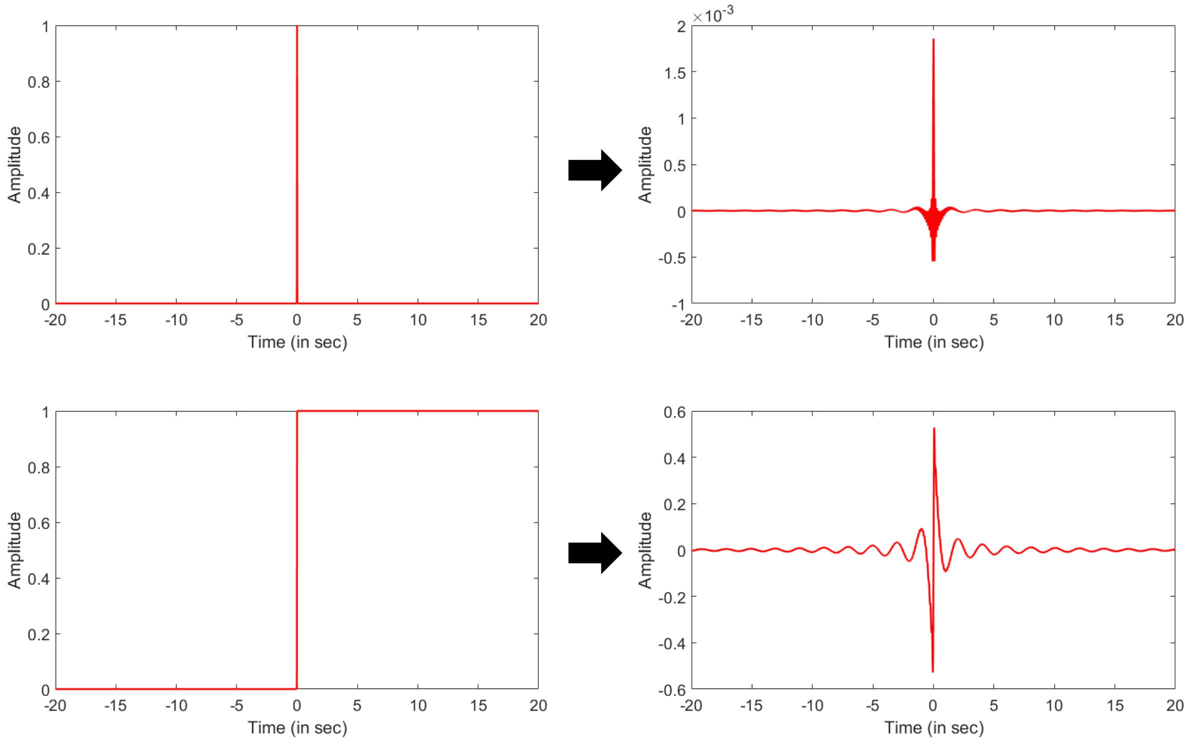


Fig. Ap11 - Impulse (upper) and step (lower) responses of the entire pipeline.

| Segment 1 (Expert 1) | | | | | |
|---|---|---|---|---|---|
| Phoneme Aligner | | | Expert labelling | | |
| Start | Stop | Phoneme | Start | Stop | Phoneme |
| 0.82045 | 0.88045 | _j | 0.8404 | 0.8768 | _j |
| 0.88045 | 0.95045 | _@ | 0.8768 | 0.9657 | _@ |
| 0.95045 | 1.0005 | _r | 0.9657 | 0.991 | _r |
| 1.0005 | 1.1005 | _u | 0.991 | 1.103 | _u |
| 1.1005 | 1.1905 | _n | 1.103 | 1.19 | _n |
| 1.1905 | 1.2705 | _w | 1.19 | 1.262 | _w |
| 1.2705 | 1.3405 | _l | 1.262 | 1.335 | _l |
| 1.3405 | 1.3905 | _n | 1.335 | 1.415 | _n |
| 1.3905 | 1.4605 | _t | 1.415 | 1.449 | _t |
| 1.4605 | 1.5705 | _v | 1.449 | 1.594 | _v |
| 1.5705 | 1.7505 | _E | 1.594 | 1.77 | _E |
| 1.7505 | 1.8805 | _f | 1.77 | 1.898 | _f |
| 1.8805 | 1.9605 | _r | 1.898 | 2.00 | _r |

| Segment 2 (Expert 1) | | | | | |
|---|---|---|---|---|---|
| Phoneme Aligner | | | Expert labelling | | |
| Start | Stop | Phoneme | Start | Stop | Phoneme |
| 1.0537 | 1.1937 | _j | 1.073 | 1.174 | _j |
| 1.1937 | 1.3137 | _o | 1.174 | 1.321 | _o |
| 1.3137 | 1.3637 | _h | 1.321 | 1.369 | _h |
| 1.3637 | 1.4637 | _A | 1.369 | 1.473 | _A |
| 1.4637 | 1.5037 | _n | 1.473 | 1.524 | _n |
| 1.5037 | 1.6137 | _m | 1.524 | 1.576 | _m |
| 1.6137 | 1.7237 | _a | 1.576 | 1.75 | _a |
| 1.7237 | 1.8237 | _k | 1.75 | 1.83 | _k |
| 1.8237 | 1.8937 | _t | 1.83 | 1.915 | _t |
| 1.8937 | 1.9737 | _v | 1.915 | 2.00 | _v |

| Segment 3 (Expert 1) | | | | | |
|---|---|---|---|---|---|
| Phoneme Aligner | | | Expert labelling | | |
| Start | Stop | Phoneme | Start | Stop | Phoneme |
| 1.0055 | 1.1255 | _s | 1.022 | 1.132 | _s |
| 1.1255 | 1.1855 | _o | 1.132 | 1.226 | _o |
| 1.1855 | 1.3255 | _f | 1.226 | 1.347 | _f |
| 1.3255 | 1.4155 | _i | 1.347 | 1.478 | _i |
| 1.5355 | 1.6155 | _k | 1.478 | 1.641 | _k |
| 1.6155 | 1.7055 | _i | 1.641 | 1.737 | _i |
| 1.7055 | 1.8255 | _s | 1.737 | 1.849 | _s |
| 1.8255 | 1.8855 | _t | 1.849 | 1.921 | _t |

| Segment 4 (Expert 1) | | | | | |
|---|---|---|---|---|---|
| Phoneme Aligner | | | Expert labelling | | |
| Start | Stop | Phoneme | Start | Stop | Phoneme |
| 0.051587 | 0.14159 | _i | 0.0732 | 0.1656 | _i |
| 0.14159 | 0.30159 | _s | 0.1656 | 0.3307 | _s |
| 0.30159 | 0.34159 | _t | 0.3307 | 0.382 | _t |
| 0.34159 | 0.38159 | _t | 0.382 | 0.4048 | _t |
| 0.38159 | 0.44159 | _w | 0.4048 | 0.4425 | _w |
| 0.44159 | 0.55159 | _a | 0.4425 | 0.5491 | _a |
| 0.55159 | 0.61159 | _l | 0.5491 | 0.6322 | _l |
| 0.61159 | 0.69159 | _f | 0.6322 | 0.7284 | _f |
| 0.72159 | 0.78159 | _p | 0.7284 | 0.7886 | _p |
| 0.78159 | 0.91159 | _a | 0.7886 | 0.9463 | _a |
| 0.91159 | 0.97159 | _r | 0.9463 | 0.9671 | _r |
| 0.97159 | 1.0816 | _s | 0.9671 | 1.104 | _s |
| 1.0816 | 1.1316 | _@ | 1.104 | 1.145 | _@ |
| 1.1616 | 1.2316 | _k | 1.145 | 1.248 | _k |
| 1.2316 | 1.3616 | _A | 1.248 | 1.396 | _A |
| 1.3616 | 1.4216 | _w | 1.396 | 1.44 | _w |
| 1.4216 | 1.5716 | _s | 1.44 | 1.598 | _s |
| 1.5716 | 1.6216 | _@ | 1.598 | 1.753 | _@ |
| 1.6216 | 1.7316 | _n | 1.753 | 1.797 | _n |

| Segment 5 (Expert 1) | | | | | |
| --- | --- | --- | --- | --- | --- |
| Phoneme Aligner | | | Expert labelling | | |
| **Start** | **Stop** | **Phoneme** | **Start** | **Stop** | **Phoneme** |
| 1.0195 | 1.0995 | _j | 1.04 | 1.09 | _j |
| 1.0995 | 1.1495 | _@ | 1.09 | 1.174 | _@ |
| 1.1495 | 1.2095 | _r | 1.174 | 1.201 | _r |
| 1.2095 | 1.2895 | _u | 1.201 | 1.315 | _u |
| 1.2895 | 1.3895 | _n | 1.315 | 1.388 | _n |
| 1.3895 | 1.4695 | _m | 1.388 | 1.489 | _m |
| 1.4695 | 1.5795 | _a | 1.489 | 1.607 | _a |
| 1.5795 | 1.6995 | _k | 1.607 | 1.709 | _k |
| 1.6995 | 1.7595 | _t | 1.709 | NA | _t |
| 1.7595 | 1.7995 | _t | NA | 1.823 | _t |
| 1.7995 | 1.8895 | _i | 1.823 | 1.875 | _i |

| Segment 6 (Expert 1) | | | | | |
| --- | --- | --- | --- | --- | --- |
| Phoneme Aligner | | | Expert labelling | | |
| **Start** | **Stop** | **Phoneme** | **Start** | **Stop** | **Phoneme** |
| 0.38587 | 0.49587 | _j | 0.4076 | 0.48 | _j |
| 0.49587 | 0.58587 | _a | 0.48 | 0.6211 | _a |
| 0.58587 | 0.68587 | _k | 0.6211 | 0.7145 | _k |
| 0.68587 | 0.75587 | _O | 0.7145 | 0.7811 | _O |
| 0.75587 | 0.78587 | _p | 0.7811 | 0.8935 | _p |
| 0.83587 | 0.92587 | _l | 0.8935 | 0.9351 | _l |
| 0.92587 | 1.0059 | _e | 0.9351 | 1.009 | _e |
| 1.0059 | 1.0559 | _n | 1.009 | 1.096 | _n |
| 1.0559 | 1.1359 | _t | 1.096 | 1.145 | _t |
| 1.1359 | 1.2259 | _v | 1.145 | 1.249 | _v |
| 1.2259 | 1.4059 | _E | 1.249 | 1.426 | _E |
| 1.4059 | 1.5159 | _f | 1.426 | 1.541 | _f |
| 1.5159 | 1.5759 | _G | 1.541 | 1.595 | _G |
| 1.5759 | 1.6259 | _r | 1.595 | 1.629 | _r |
| 1.6259 | 1.7659 | _E | 1.629 | 1.804 | _E |
| 1.7659 | 1.8459 | _z | 1.804 | 1.858 | _z |
| 1.8459 | 1.8859 | _@ | 1.858 | 1.914 | _@ |

| Segment 7 (Expert 2) | | | | | |
| --- | --- | --- | --- | --- | --- |
| Phoneme Aligner | | | Expert labelling | | |
| **Start** | **Stop** | **Phoneme** | **Start** | **Stop** | **Phoneme** |
| 0.23562 | 0.32562 | _j | 0.237 | 0.311 | _j |
| 0.32562 | 0.41562 | _a | 0.311 | 0.439 | _a |
| 0.41562 | 0.52562 | _k | 0.439 | 0.523 | _k |
| 0.52562 | 0.58562 | _O | 0.523 | 0.582 | _O |
| 0.58562 | 0.65562 | _p | 0.582 | 0.68 | _p |
| 0.65562 | 0.76562 | _z | 0.68 | 0.748 | _z |
| 0.76562 | 0.80562 | _u | 0.748 | 0.822 | _u |
| 0.80562 | 0.92562 | _k | 0.822 | 0.933 | _k |
| 0.92562 | 1.0356 | _t | 0.933 | 1.044 | _t |
| 1.0356 | 1.0756 | _w | 1.044 | 1.0755 | _w |
| 1.0756 | 1.1756 | _a | 1.075 | 1.175 | _a |
| 1.1756 | 1.2556 | _l | 1.175 | 1.268 | _l |
| 1.2556 | 1.3556 | _f | 1.268 | 1.354 | _f |
| 1.3556 | 1.4356 | _G | 1.354 | 1.434 | _G |
| 1.4356 | 1.4756 | _r | 1.434 | 1.47 | _r |
| 1.4756 | 1.5356 | _u | 1.47 | 1.549 | _u |
| 1.5356 | 1.6056 | _n | 1.549 | 1.641 | _n |
| 1.6056 | 1.6756 | _@ | 1.641 | 1.704 | _@ |
| 1.6756 | 1.7456 | _b | 1.704 | 1.752 | _b |
| 1.7456 | 1.8856 | _o | 1.752 | 1.9 | _o |

| Segment 8 (Expert 2) | | | | | |
|---|---|---|---|---|---|
| Phoneme Aligner | | | Expert labelling | | |
| **Start** | **Stop** | **Phoneme** | **Start** | **Stop** | **Phoneme** |
| 0.0038322 | 0.16383 | _a | 0.005 | 0.172 | _a |
| 0.16383 | 0.20383 | _r | 0.172 | 0.195 | _r |
| 0.20383 | 0.31383 | _a | 0.195 | 0.322 | _a |
| 0.31383 | 0.42383 | _z | 0.322 | 0.429 | _z |
| 0.42383 | 0.47383 | _u | 0.429 | 0.488 | _u |
| 0.47383 | 0.60383 | _k | 0.488 | 0.601 | _k |
| 0.60383 | 0.72383 | _t | 0.601 | 0.728 | _t |
| 0.72383 | 0.81383 | _i | 0.728 | 0.819 | _i |
| 0.81383 | 0.94383 | _n | 0.819 | 0.948 | _n |
| 0.94383 | 1.0338 | _G | 0.948 | 1.044 | _G |
| 1.0338 | 1.1438 | _e | 1.044 | 1.143 | _e |
| 1.1438 | 1.1938 | _l | 1.143 | 1.207 | _l |
| 1.1938 | 1.2638 | _@ | 1.207 | 1.258 | _@ |
| 1.3038 | 1.3538 | _p | 1.258 | 1.373 | _p |
| 1.3538 | 1.4338 | _E | 1.373 | 1.436 | _E |
| 1.4338 | 1.4838 | _n | 1.436 | 1.487 | _n |
| 1.4838 | 1.5538 | _@ | 1.487 | 1.556 | _@ |
| 1.5538 | 1.6538 | _n | 1.556 | 1.697 | _n |

| Segment 9 (Expert 2) | | | | | |
|---|---|---|---|---|---|
| Phoneme Aligner | | | Expert labelling | | |
| **Start** | **Stop** | **Phoneme** | **Start** | **Stop** | **Phoneme** |
| 0.36626 | 0.41626 | _d | 0.366 | 0.418 | _d |
| 0.41626 | 0.57626 | _a | 0.418 | 0.593 | _a |
| 0.57626 | 0.66626 | _v | 0.593 | 0.676 | _v |
| 0.66626 | 0.74626 | _l | 0.676 | 0.742 | _l |
| 0.74626 | 0.80626 | _t | 0.742 | 0.827 | _t |
| 0.80626 | 0.90626 | _z | 0.827 | 0.888 | _z |
| 0.90626 | 0.94626 | _u | 0.888 | 0.956 | _u |
| 0.94626 | 1.0463 | _k | 0.956 | 1.043 | _k |
| 1.0463 | 1.1263 | _t | 1.043 | 1.113 | _t |
| 1.1263 | 1.2363 | _v | 1.113 | 1.247 | _v |
| 1.2363 | 1.3963 | _i | 1.247 | 1.398 | _i |
| 1.3963 | 1.4563 | _r | 1.398 | 1.421 | _r |
| 1.4563 | 1.5163 | _w | 1.421 | 1.469 | _w |
| 1.5163 | 1.5663 | _l | 1.469 | 1.582 | _l |
| 1.5663 | 1.6463 | _t | 1.582 | 1.651 | _t |
| 1.6463 | 1.7163 | _@ | 1.651 | 1.709 | _@ |
| 1.7163 | 1.7963 | _d | 1.709 | 1.808 | _p |
| 1.7963 | 1.8863 | _E | 1.808 | 1.878 | _E |
| 1.8863 | 1.9363 | _n | 1.878 | 1.937 | _n |
| 1.9363 | 1.9963 | _@ | 1.937 | 2 | _@ |

| Segment 10 (Expert 2) | | | | | |
|---|---|---|---|---|---|
| Phoneme Aligner | | | Expert labelling | | |
| **Start** | **Stop** | **Phoneme** | **Start** | **Stop** | **Phoneme** |
| 0.10111 | 0.17111 | _t | 0.092 | 0.168 | _t |
| 0.20111 | 0.31111 | _A | 0.224 | 0.307 | _A |
| 0.31111 | 0.40111 | _x | 0.307 | 0.396 | _x |
| 0.40111 | 0.45111 | _t | 0.396 | 0.449 | _t |
| 0.45111 | 0.53111 | _z | 0.449 | 0.54 | _z |
| 0.53111 | 0.57111 | _w | 0.54 | 0.571 | _w |
| 0.57111 | 0.64111 | _A | 0.571 | 0.645 | _A |
| 0.64111 | 0.72111 | _r | 0.645 | 0.727 | _r |
| 0.72111 | 0.80111 | _t | 0.727 | 0.789 | _t |
| 0.80111 | 0.83111 | _@ | 0.795 | 0.83 | _@ |
| 0.83111 | 0.91111 | _m | 0.83 | 0.903 | _m |
| 0.91111 | 1.0011 | _A | 0.903 | 0.995 | _A |
| 1.0011 | 1.1011 | _n | 0.995 | NA | _n |
| 1.1011 | 1.1311 | _d | NA | 1.128 | _d |
| 1.1311 | 1.2011 | _@ | 1.128 | 1.209 | _@ |
| 1.2011 | 1.3611 | _n | 1.209 | 1.388 | _n |

| Segment 11 (Expert 2) | | | | | |
|---|---|---|---|---|---|
| Phoneme Aligner | | | Expert labelling | | |
| **Start** | **Stop** | **Phoneme** | **Start** | **Stop** | **Phoneme** |
| 0.25007 | 0.36007 | _s | 0.266 | 0.357 | _s |
| 0.36007 | 0.40007 | _O | 0.357 | 0.419 | _O |
| 0.40007 | 0.55007 | _f | 0.419 | 0.552 | _f |
| 0.55007 | 0.68007 | _i | 0.552 | 0.704 | _i |
| 0.71007 | 0.81007 | _m | 0.704 | 0.808 | _m |
| 0.81007 | 0.91007 | _a | 0.808 | 0.931 | _a |
| 0.91007 | 1.0101 | _k | 0.931 | 1.014 | _k |
| 1.0101 | 1.0801 | _t | 1.014 | 1.111 | _t |
| 1.0801 | 1.1801 | _v | 1.111 | 1.187 | _v |
| 1.1801 | 1.3701 | _E+ | 1.187 | 1.373 | _E+ |
| 1.3701 | 1.4501 | _f | 1.373 | 1.414 | _f |
| 1.4801 | 1.5201 | _b | 1.414 | 1.527 | _b |
| 1.5201 | 1.5701 | _r | 1.527 | 1.564 | _r |
| 1.5701 | 1.7201 | _Y+ | 1.564 | 1.717 | _Y+ |
| 1.7201 | 1.7601 | _n | 1.717 | 1.777 | _n |
| 1.7601 | 1.8201 | _@ | 1.777 | 1.83 | _@ |
| 1.8601 | 1.9101 | _p | 1.83 | 1.929 | _p |

Table Ap1 – Evaluation of the automatic phoneme alignment by two audiology experts. Both experts randomly selected six 2s-segments in the different Matrix Speech materials and manually labelled them. In columns 1 to 3, the different phonemes with their corresponding starting and stopping point within the 2s-segment using the automatic phoneme aligner. In columns 4 to 6, the corresponding expert's manual labelling.

| Segment 12 (Expert 2) | | | | | |
|---|---|---|---|---|---|
| Phoneme Aligner | | | Expert labelling | | |
| **Start** | **Stop** | **Phoneme** | **Start** | **Stop** | **Phoneme** |
| 0.38871 | 0.47871 | _d | 0.371 | 0.475 | _d |
| 0.47871 | 0.62871 | _a | 0.475 | 0.652 | _a |
| 0.62871 | 0.70871 | _v | 0.652 | 0.719 | _v |
| 0.70871 | 0.77871 | _I | 0.719 | 0.793 | _I |
| 0.77871 | 0.89871 | _t | 0.739 | 0.889 | _t |
| 0.92871 | 0.97871 | _m | 0.932 | 0.988 | _m |
| 0.97871 | 1.0987 | _a | 0.988 | 1.12 | _a |
| 1.0987 | 1.2087 | _k | 1.12 | 1.215 | _k |
| 1.2087 | 1.2587 | _t | 1.215 | 1.247 | _t |
| 1.2587 | 1.2987 | _t | 1.247 | 1.316 | _t |
| 1.2987 | 1.4287 | _i | 1.316 | 1.389 | _i |
| 1.4287 | 1.5487 | _n | 1.389 | NA | _n |
| 1.5487 | 1.5887 | _b | NA | 1.592 | _b |
| 1.5887 | 1.6287 | _l | 1.592 | 1.64 | _l |
| 1.6287 | 1.7587 | _A+ | 1.64 | 1.732 | _A+ |
| 1.7587 | 1.7887 | _w | 1.732 | 1.807 | _w |
| 1.7887 | 1.8387 | _@ | 1.807 | 1.84 | _@ |
| 1.8387 | 1.9087 | _m | 1.84 | 1.906 | _m |
| 1.9087 | 1.9987 | _A | 1.906 | 2 | _A |
